# An exploration of Prevotella-rich microbiomes in HIV and men who have sex with men

**DOI:** 10.1101/424291

**Authors:** Abigail JS Armstrong, Michael Shaffer, Nichole M Nusbacher, Christine Griesmer, Suzanne Fiorillo, Jennifer M Schneider, C Preston Neff, Sam X Li, Andrew P Fontenot, Thomas Campbell, Brent E Palmer, Catherine A Lozupone

## Abstract

**Background:** Gut microbiome characteristics associated with HIV infection are of intense research interest but a deep understanding has been challenged by confounding factors across studied populations. Notably, a Prevotella-rich microbiome described in HIV-infected populations is now understood to be common in men who have sex with men (MSM) regardless of HIV status, but driving factors and potential health implications are unknown.

**Results:** Here we further define the MSM-associated gut microbiome and describe compositional differences between the fecal microbiomes of Prevotella-rich MSM and non-MSM that may underlie observed pro-inflammatory properties. Furthermore, we show relatively subtle gut microbiome changes in HIV infection in MSM and women that include an increase in potential pathogens that is ameliorated with antiretroviral therapy (ART). Lastly, using a longitudinal cohort, we describe microbiome changes that happen after ART initiation.

**Conclusions:** This study provides an in-depth characterization of microbiome differences that occur in a US population infected with HIV and demonstrates the degree to which these differences may be driven by lifestyle factors, ART and HIV infection itself. Understanding microbiome compositions that occur with sexual behaviors that are high-risk for acquiring HIV and untreated and ART-treated HIV infection will guide the investigation of immune and metabolic functional implications to ultimately target the microbiome therapeutically.

## Background

One bodily fluid people rarely think of as related to human immunodeficiency virus (HIV) infection is feces. However, the gut microbiome may hold the key for preventing disease transmission and progression and for improving the quality of life for the 1.2 million Americans and 36.7 million individuals worldwide with chronic HIV infection for several reasons (1). First, the gut microbiota can induce T cell activation (2) and since HIV preferentially infects activated T cells, microbiome composition at mucosal sites, specifically in the rectum/anus, may influence disease transmission in men who have sex with men (MSM) (3–5). Second, increased peripheral immune activation driven by translocation of gut bacterial components has been linked to HIV disease progression in both untreated and treated infection (6), and the high peripheral immune activation may be influenced by gut microbiota with more pro-inflammatory components (2). Finally, although life expectancy of individuals infected with HIV has been greatly increased by antiretroviral therapy (ART) (7), there has been a concurrent increase in non-infectious comorbidities including metabolic and cardiovascular diseases associated with chronic inflammation (7–9). Outside the context of HIV infection, these diseases have been associated with microbiome dysbiosis (10–12).

Understanding compositional changes of the gut microbiome the occur with HIV infection is hindered by the complexity of the HIV-infected population and the myriad confounding factors (13). Most notably, a prominent HIV-associated microbiome alteration observed by our group (14) and independently by several others (15–17) was an increase in the genus *Prevotella* and decrease in the genus *Bacteroides*. However, recent studies have shown that MSM have a Prevotella-rich fecal (2, 18) and rectal mucosal (9) microbiome regardless of HIV infection status. HIV-negative MSM are therefore necessary controls for distinguishing the effects of HIV infection itself on microbiome composition, especially in US populations where the majority of HIV-positive individuals are MSM (5).

Potential health implications of Prevotella-rich microbiomes have been a topic of much interest (19). Human gut microbiomes across many studies show related clustering patterns based on their composition, leading to the suggestion that human microbiomes form distinct “enterotypes” characterized by particular community structures (20). The Prevotella enterotype, which is characterized by a high relative abundance of the genus *Prevotella*, occurs in approximately 18% of healthy individuals in western populations (20). It has been linked with diets low in animal products and high in carbohydrates/fiber (21) and with beneficial metabolic effects in the context of a high fiber diet (22). Prevotella-rich microbiomes are also typical of healthy individuals in agrarian cultures (23). However, high Prevotella has also been described in inflammatory states such as rheumatoid arthritis (24) and has been linked with obesity (25, 26) and insulin resistance (27). Furthermore, we have shown that in *in vitro* stimulations the Prevotella-rich fecal microbiome of MSM induces higher activation of human innate immune cells compared to non-MSM (2), supporting the idea that the MSM-microbiome is potentially a contributing factor to increased inflammation observed in MSM both systemically (28, 29) and in rectal mucosa (9).

In this paper, we characterize the fecal microbiome of MSM and non-MSM to explore potential driving factors and health implications of a Prevotella-rich microbiomes in MSM individuals. We also further characterize gut microbiome attributes that are associated with HIV infection and ART. Taken together, this study provides an in-depth characterization of microbiome differences that occur in a US population infected with HIV and demonstrates the degree to which these differences may be driven by lifestyle factors, ART and HIV infection itself. This understanding will help to guide efforts to investigate the functional implications of these differences and ultimately target the microbiome therapeutically.

## Results

### Cohort description

This paper compares the fecal microbiome of 217 individuals as assessed by sequencing the V4 region of the 16S ribosomal RNA (rRNA) gene (Table 1; Additional Table 1). 68% of the cohort filled out a sexual preference/behavior questionnaire (Table 2; Additional Table 2). Taken together, our analysis includes HIV-positive individuals on ART; HIV-positive, ART-naïve individuals; and a seronegative cohort. The seronegative cohort includes 35 MSM of which the majority (n=32) engaged in high risk behaviors within the prior year; 29 men who identify as heterosexual and did not report engaging in any sexual risk factors (referred to here as MSW; men who have sex with women); and 41 women, 3 of whom reported in engaging in receptive anal intercourse within a year of sample collection (Table 1, 2).

**Table 1.**
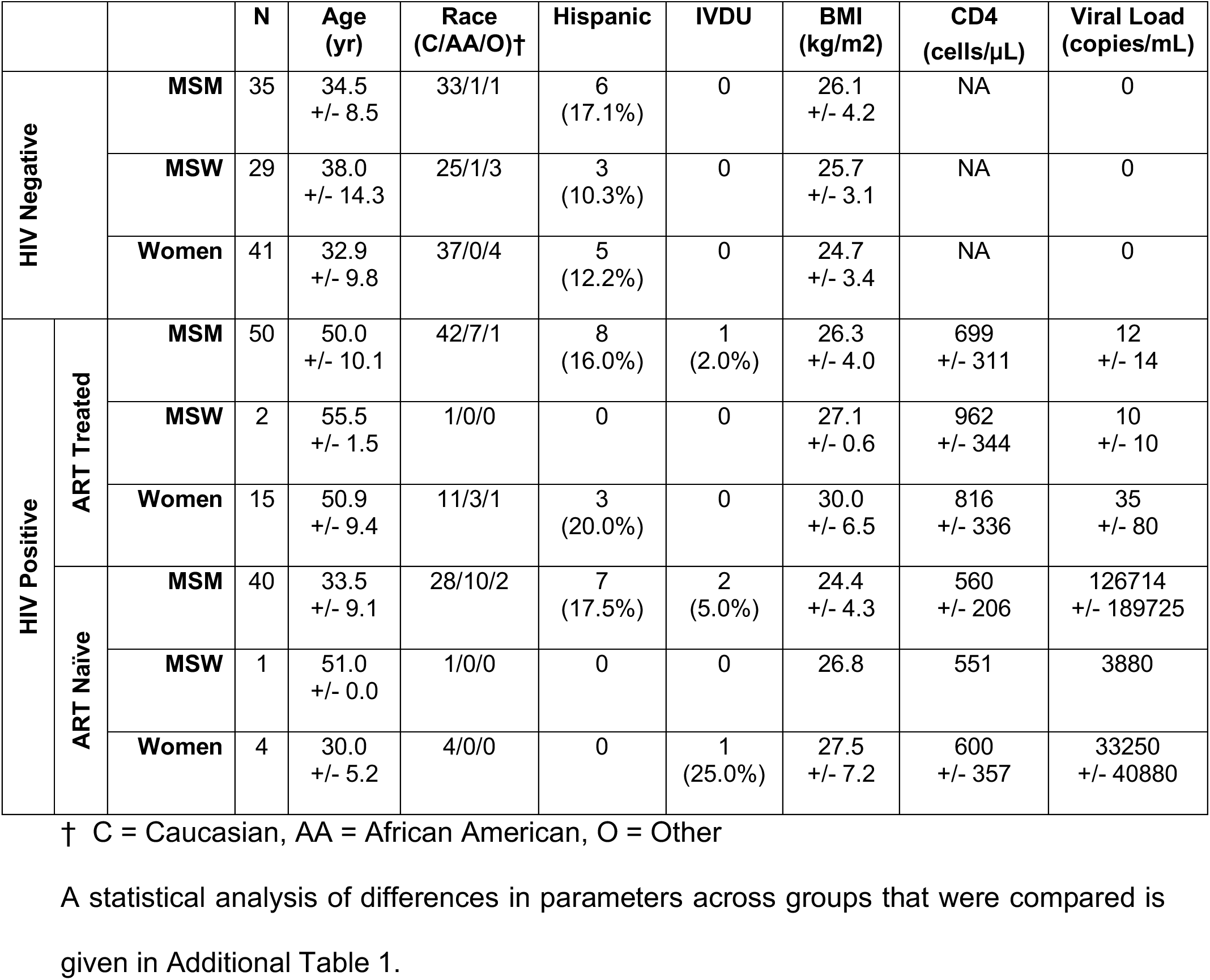
Cohort Description

**Table 2.**
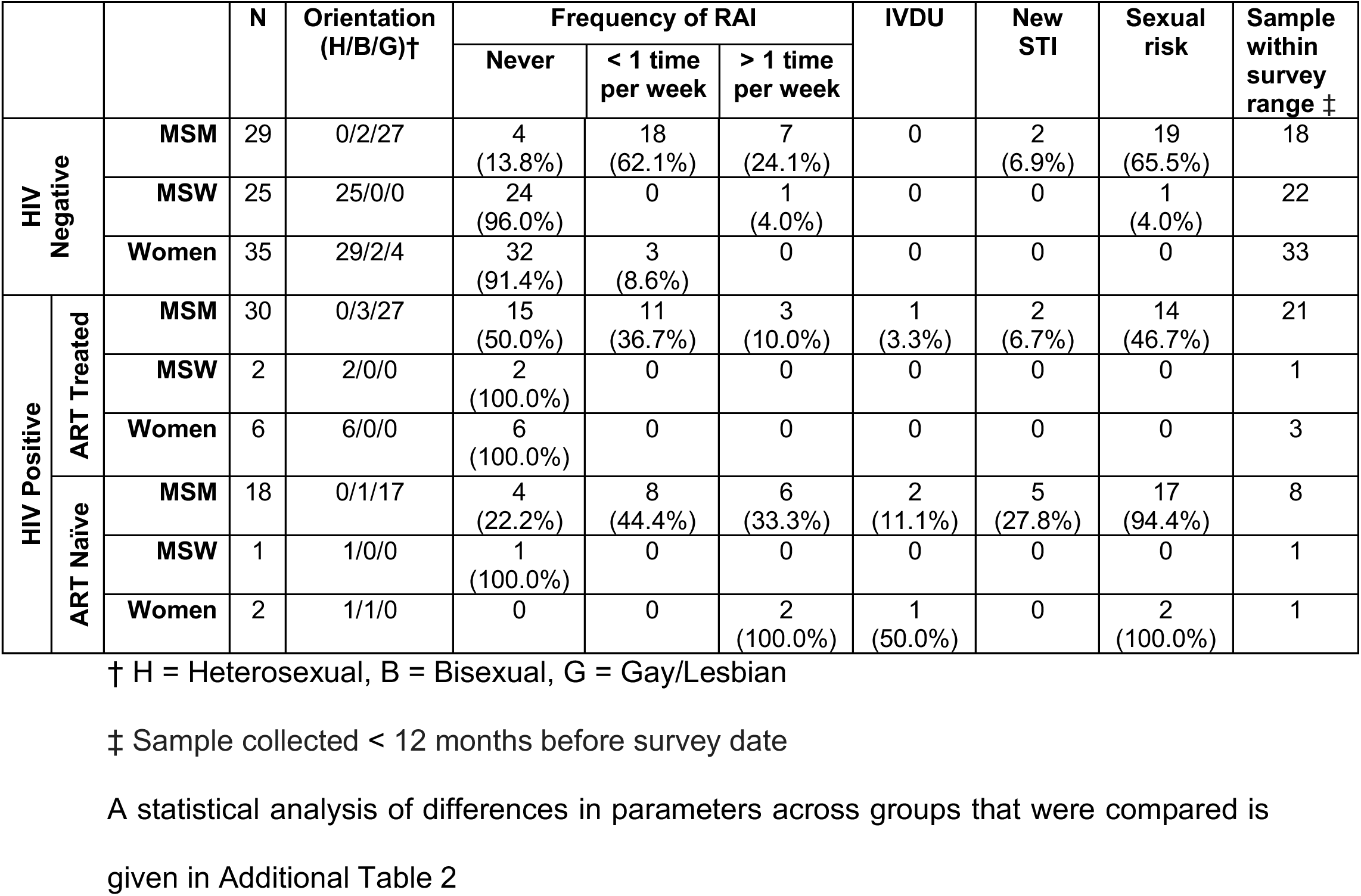
Behavioral Data

Almost all of the HIV-positive men who responded to our behavior questionnaire (95%) were sexually active MSM, usually engaging in anal intercourse with other HIV-positive individuals (Table 2). Although 25 of the 93 HIV-positive men did not respond to our behavior questionnaire, we included these 25 individuals in our HIV-positive, MSM cohort. To confirm that potential false identification as MSM would not affect our results, we also performed analysis without the 25 HIV-positive men of unsure MSM status, and there was no significant change in results (Additional Table 3). One of the HIV-positive males in the ART group is a female to male (FTM) transgender who identifies as MSW and is categorized as such in our analyses.

The ART-treated cohort all had plasma HIV RNA at or below the limits of detection. There were significantly higher CD4+ T cell numbers (cells/μL) in HIV-positive MSM on ART compared to the ART-naïve cohorts but no significant difference between ART-treated and ART-naïve HIV-positive women (p < 0.05 and p = 0.32; Kruskal-Wallis test; Additional Table 1). Of note, 12 HIV-positive individuals had a CD4+ T cell count below 300/μL, 6 in the ART cohort (1 female, 5 MSM) and 6 in the ART naïve cohort (1 female, 5 MSM). These numbers – an estimate of HIV disease progression – suggest very few individuals in our cohort with immune deficiencies consistent with advanced HIV infection.

### Sexual behavior is the strongest descriptor of compositional variation in the gut microbiome

We observed clear clustering by gender/sexual behavior but not HIV infection status when applying Principal Coordinates Analysis (PCoA) to weighted (Figure 1 A, C, Additional Figure 1B) and unweighted (Figure 1B, Additional Figure 1A) UniFrac distance matrices as well as abundance and binary Jaccard (Additional Figure 2A,B). Applying the Adonis test showed significant effects for MSM and HIV with weighted Unifrac, which explained 16.4% and 1.4% of the variation respectively, and for MSM, HIV, and ART with unweighted Unifrac, which explained 7.0%, 0.9%, 0.7% respectively (Additional Table 3).

**Figure 1.**
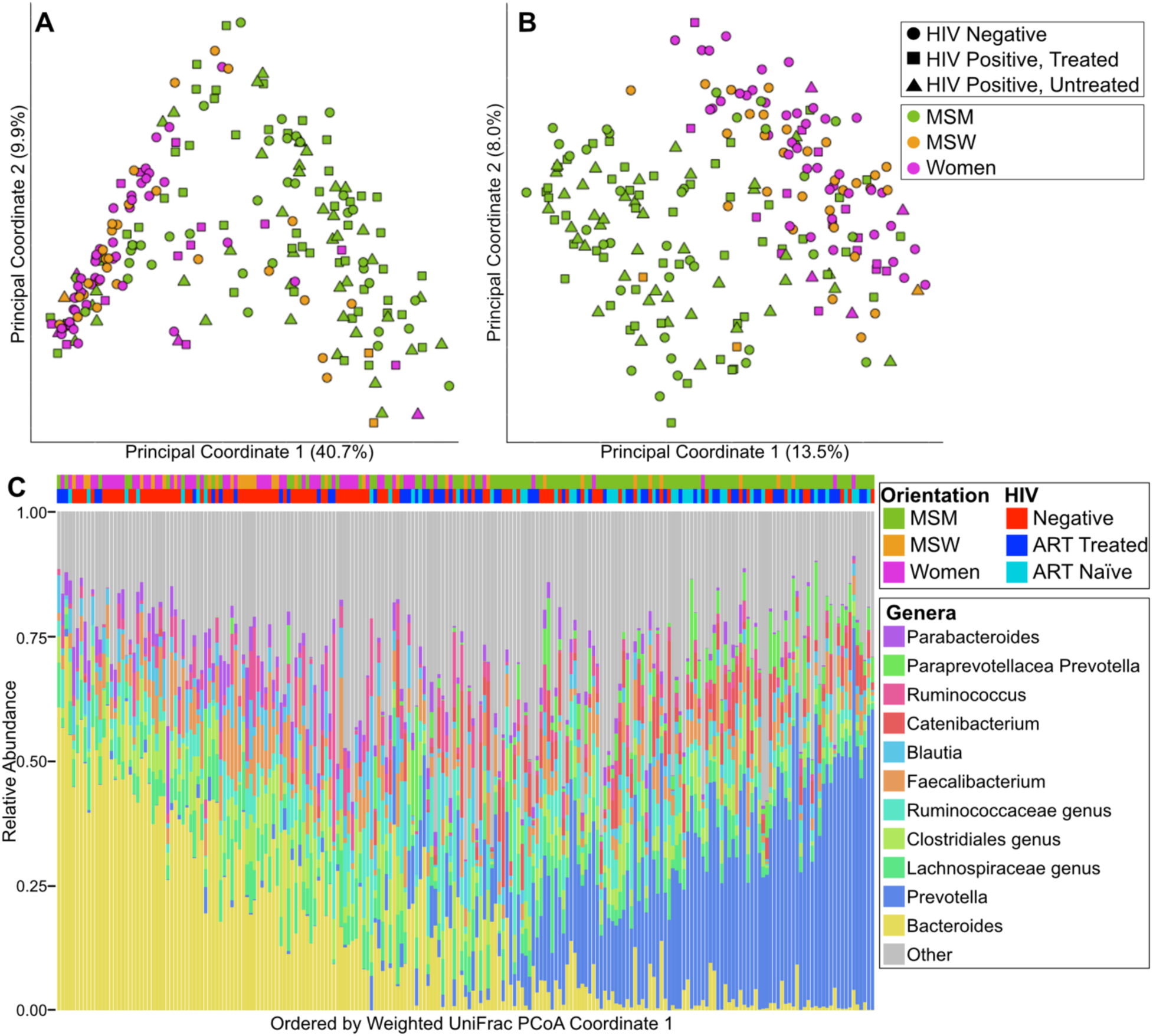
MSM is more strongly associated with Prevotella richness than HIV infection. **A.** Weighted UniFrac PCoA and **B.** Unweighted UniFrac with points colored by orientation and shaped by HIV status. **C.** Genus level taxonomic summary plot. (Bottom) Genera with mean relative less than 2% abundance are binned together into the category “Other”. Each column represents one individual. (Top) samples are marked with HIV status and orientation. Each column corresponds to the genus plot below. Samples are ordered by the coordinate on the weighted UniFrac principle coordinate 1.

In order to empirically determine any clustering within the data, we performed the standard “enterotyping” methods as previously described (21). In short, clusters are defined on a Jensen-Shannon Divergence matrix by partition around medoids clustering (Additional Figure 3B). Calculating the Silhouette index on our data revealed optimum clustering into two clusters (Additional Figure 3A). These cluster are primarily defined by a dominance of the genus *Bacteroides* or *Prevotella* (Additional Figure 3C). We subsequently refer to individuals in the *Prevotella* dominant enterotype as individuals with Prevotella-rich microbiomes. MSM were more likely to be Prevotella-rich than females and MSW (p < 0.001; Fisher’s Exact Test).

HIV-negative and HIV-positive MSM had significantly greater Phylogenetic Diversity (PD; (30)) and observed Operational Taxonomic Units (OTUs, defined at a 99% identity threshold) compared to non-MSM (p < 0.05; Kruskal-Wallis test; Figure 2A) and many significantly different OTUs (Additional table 4). The most striking difference in MSM compared to non-MSM is the increase in relative abundance of the *Prevotella* genus. On average HIV-negative MSM had 3.9 times higher relative abundance of *Prevotella* compared to HIV-negative non-MSM.

**Figure 2.**
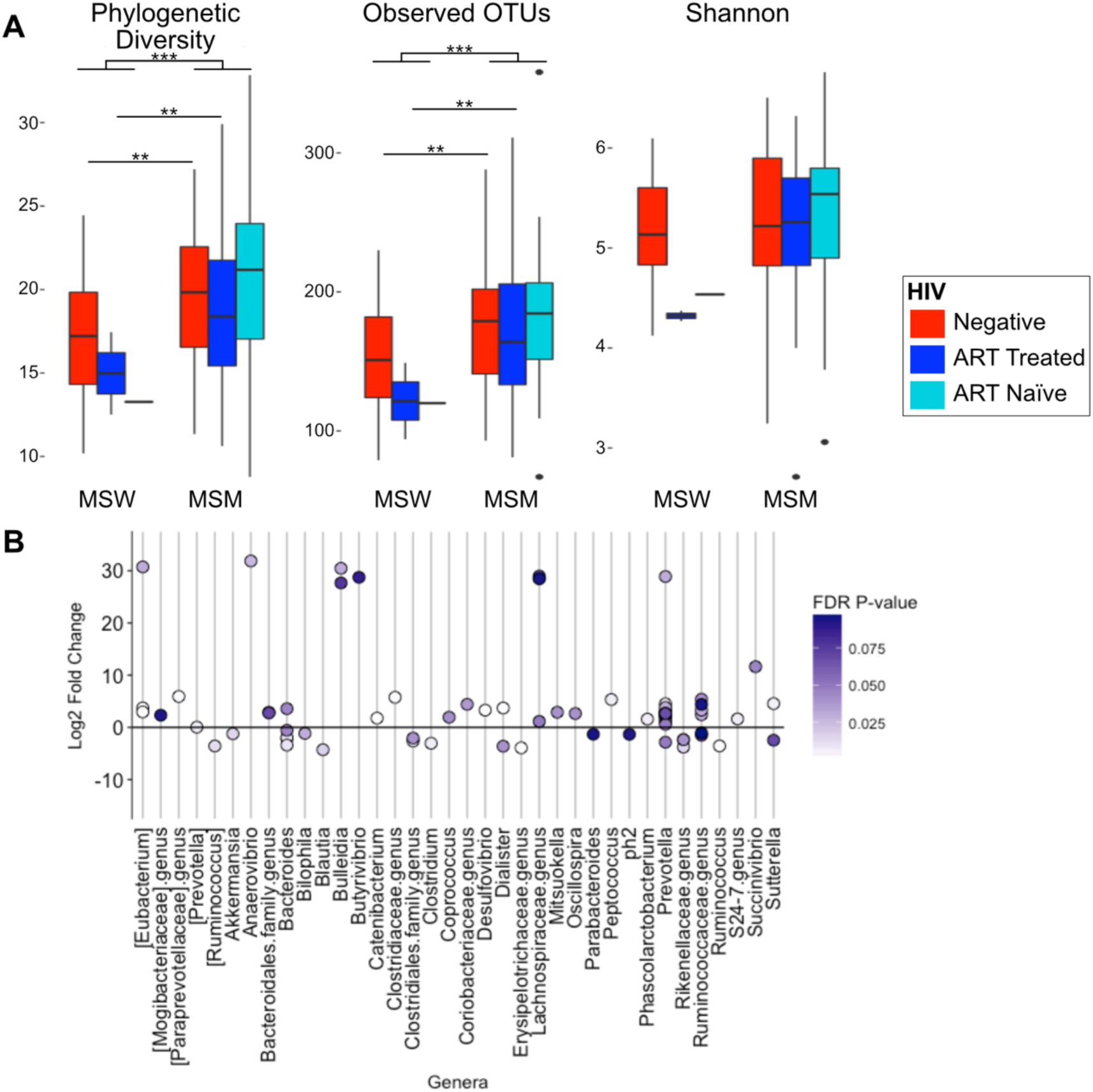
HIV negative MSM have significantly different microbiome composition compared to HIV-negative MSW. **A.** MSM have significantly higher alpha diversity than MSW with and without controlling for HIV status. (Kruskal-Wallis test; *p<0.05, **p<0.01, ***p<0.001, NS=not significant) **B.** Several OTUs are significantly different between HIV-negative Prevotella-rich MSM and HIV-negative Prevotella-rich non-MSM. The significant (Kruskal-Wallis; FDR-corrected p < 0.1) OTUs are displayed as the Log2 abundance fold change in MSM compared to non-MSM. More detailed information on the significant OTUs is given in Additional Table 6.

### Prevotella-rich microbiomes differ in MSM compared to non-MSM

Prevotella-rich non-MSM clustered together with Prevotella-rich MSM in weighted but not unweighted UniFrac (Figure 1C; Additional Figure 1A), indicating compositional differences in rare taxa. To understand these enteroyptes in the context of other studies, our samples were classified using enterotypes.org (31), which uses large-scale microbiome studies including the Human Microbiome Project (32) and MetaHIT (33) to form an enterotype reference space to which other data can be compared. We found that the proportion of Prevotella-rich individuals as defined by “enterotyping” in our study who did not fit within the general Prevotella-rich reference space defined in enterotypes.org was significantly higher for MSM (81.4% of individuals) than non-MSM (46.7%) (Additional Table 5; Fisher’s test; p < 0.05). Prevotella-rich, HIV-negative MSM (n=25) and non-MSM (n=9) had significantly different relative abundances of 39 OTUs in 21 different genera (Kruskal-Wallis test; FDR P < 0.1; Figure 2B. Additional Table 6). Interestingly the OTUs with the largest increase and decrease in MSM were both identified as *Prevotella copri*, suggesting strain-level differences in *Prevotella* (Additional Table 6).

To understand the highly interactive set of co-occurring microbes within these Prevotella-rich microbiomes, we used SparCC correlations to build a co-occurrence network of OTU-OTU correlations in Prevotella-rich, HIV-negative MSM (Additional Figure 4; Additional Table 7) and separately in Prevotella-rich, HIV-negative non-MSM (Additional Table 8). We had a deep sampling of the Prevotella-rich, HIV-negative MSM (n=25) compared to the non-MSM (n = 9) and thus designed our analysis to determine the degree to which OTU-OTU relationships that were reliably observed in the MSM group at a sample depth of 9 were observed in the non-MSM at the same depth. Specifically, we performed jackknifing at an n of 9 on the MSM samples 100 times and weighted the edges in the resulting network by the fraction of times the edge was observed. We then evaluated whether high-confidence edges (correlations) in the MSM network (weight of 75% or greater) were found in the non-MSM network. Only 18.6% of high-confidence edges (n = 18) in the MSM network were shared with the non-MSM network. Of note 8.2% (n = 8) of the high-confidence edges in the MSM network have one or both of the nodes within the Prevotella genus; however, none of these are shared with the non-MSM network.

### Potential drivers of Prevotella-richness in MSM

We collected behavioral data including frequency of receptive anal intercourse (RAI) (Table 2) on a total of 77 MSM individuals. As some subjects received the questionnaire retrospectively, our analysis included only the 47 MSM individuals who answered within the one year timeframe of sample collection that the behavior questions referred to (Table 2). However, we did not find any significant association with microbiome taxonomy or alpha or beta diversity between MSM who engaged in RAI (n = 31) and those who did not (n = 15) when controlling for sexual orientation and HIV status (Additional Figure 5; taxonomy and alpha diversity – Kruskal-Wallis test; beta diversity – adonis test; p > 0.05). We also did not find a significant association with RAI frequency (<1 time per week compared to >1 time per week). The 5 women (3 HIV negative and 2 HIV positive) who reported engaging in RAI – 2 at relatively high frequency (>1 time per week) – were all Bacteroides rich and clustered with non-RAI-engaging women and MSW in Unifrac PCoA (Additional Figure 5).

Because diet composition has been associated with enterotypes (21), we also collected diet information on a subset of the subjects (n = 98). Comparison of diets between HIV-negative MSM (n = 24) and non-MSM (n = 45) revealed several significant differences in diet composition in both data normalized to 1000kcal (Additional Table 9) and non-normalized data (Additional Table 10) (Mann-Whitney U; FDR p < 0.1). Most notably, MSM reported eating more ounces of lean meat from beef, pork, veal, lamb, and game and fewer servings of fruit and grams of dietary fiber (Figure 3). However, these diet changes do not appear to be driving our reported differences between MSM and MSW/females as linear models that included diet did not change the significance of the top ten most significantly different OTUs with MSM status (Additional Table 11). Lastly, though previous studies have suggested an effect of IV drug usage on microbiome composition in HIV (34), our cohort did not include a sufficient number of IV drug users (n = 4) to infer any statistically significant influence.

**Figure 3.**
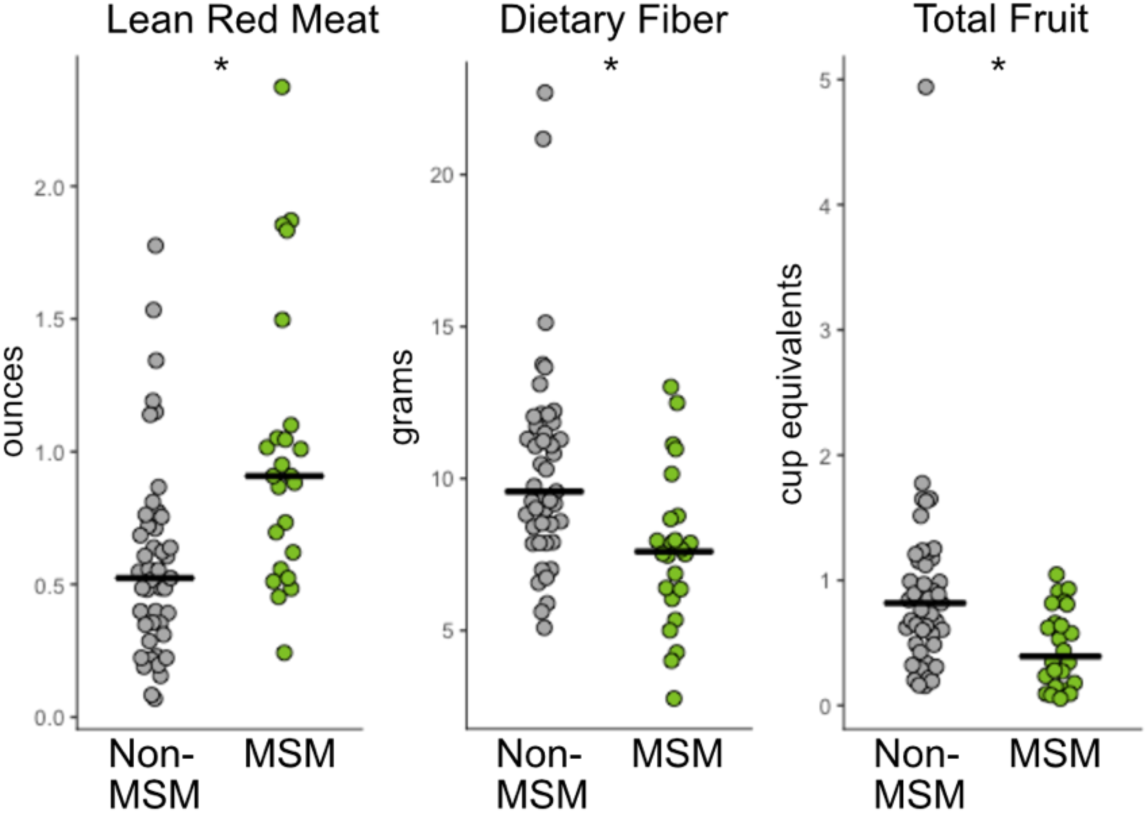
FFQ diet analysis of MSM compared to non-MSM shows MSM eat significantly few servings of red meat and fruits and less fiber. Reported values are normalized per 1000kcals. More detail on significant diet components is given in Additional Tables 9 and 10 (Kruskal-Wallis test;; *p<0.05)

### Identifying HIV-associated microbiome differences while controlling for sexual behavior and gender

We stratified our cohort into analyses containing MSM only and women only to investigate changes in the microbiome that may occur with HIV infection and ART (Figure 4A). We observed no significant difference in alpha diversity with HIV infection status in MSM (Fig. 2). In MSM, six OTUs were significantly different between HIV-negative, HIV-positive, untreated, and HIV-positive on ART based on Kruskal-Wallis with the Dunn’s post hoc test and FDR correction (Table 3; Additional Table 12). The relative abundances of these OTUs followed one of two trends: First, no significant difference in untreated HIV infection compared to HIV-negative controls and significant decrease in treated HIV-infection compared to untreated. This trend suggests microbes not altered due to HIV infection but depleted with ART and includes an OTU identified as *Fenollaria massiliensis* and an OTU within the *Peptoniphilus* genus. Second, a significant increase in abundance in untreated HIV infection compared to HIV-negative controls and a significant decrease in treated compared to untreated HIV infection. This trend suggests microbes that may respond to viral load and immune depletion in untreated infection that is corrected by ART and includes OTUs identified as *Fusobacterium equinum*, *Turicibacter sanguinis*, *Finegoldia magna*, and *Streptococcus* sp. (*S. mitis* group).

**Figure 4.**
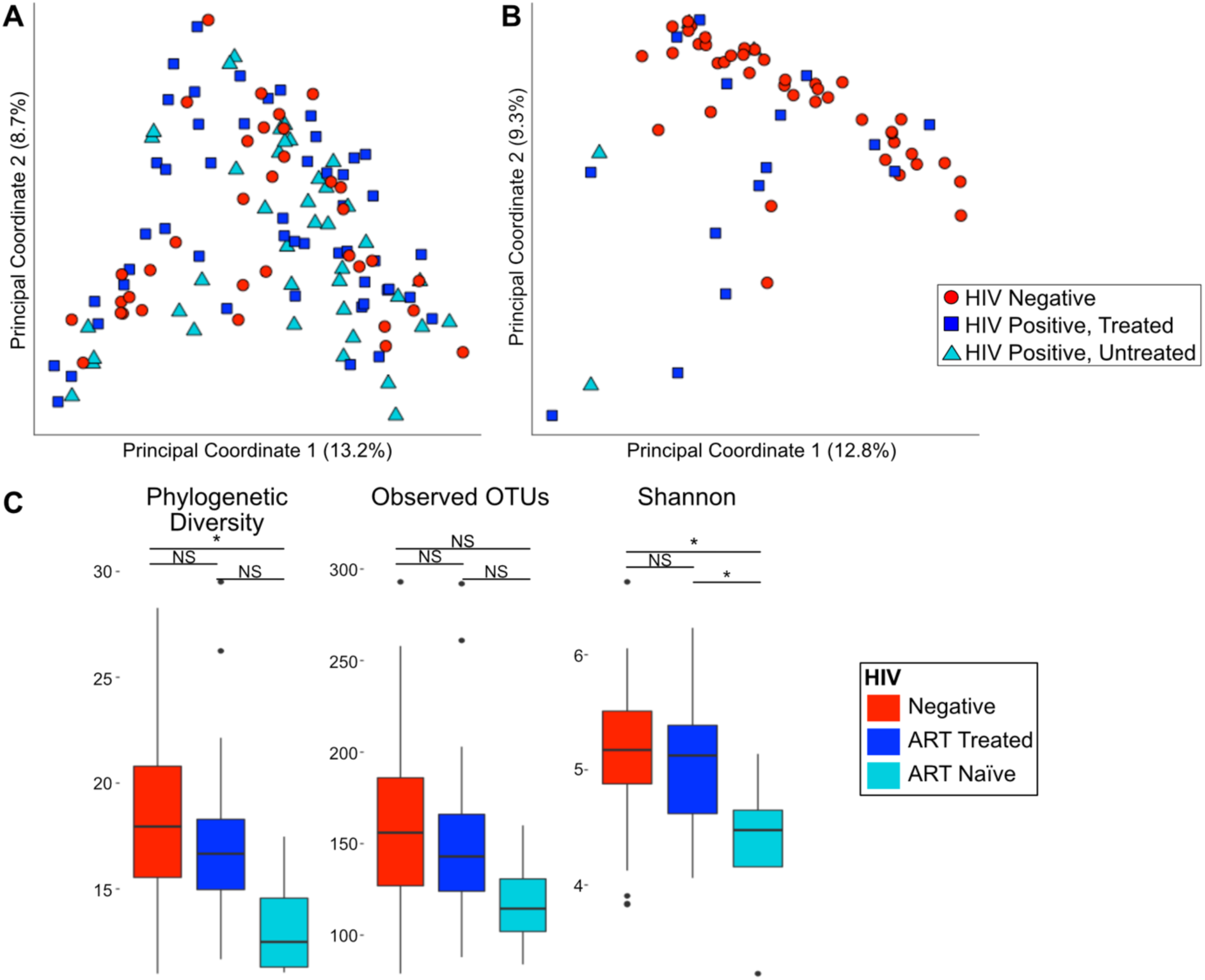
HIV specific changes in Women in MSM. Weighted UniFrac PCoA of **A.** MSM only and **B.** women only shows that only half of HIV-positive women cluster apart from HIV-negative women whereas there is no distinctive clustering of HIV in MSM. **C.** Alpha diversity differences in women only (Kruskal-Wallis test; *p<0.05, NS=not significant)

**Table 3.**
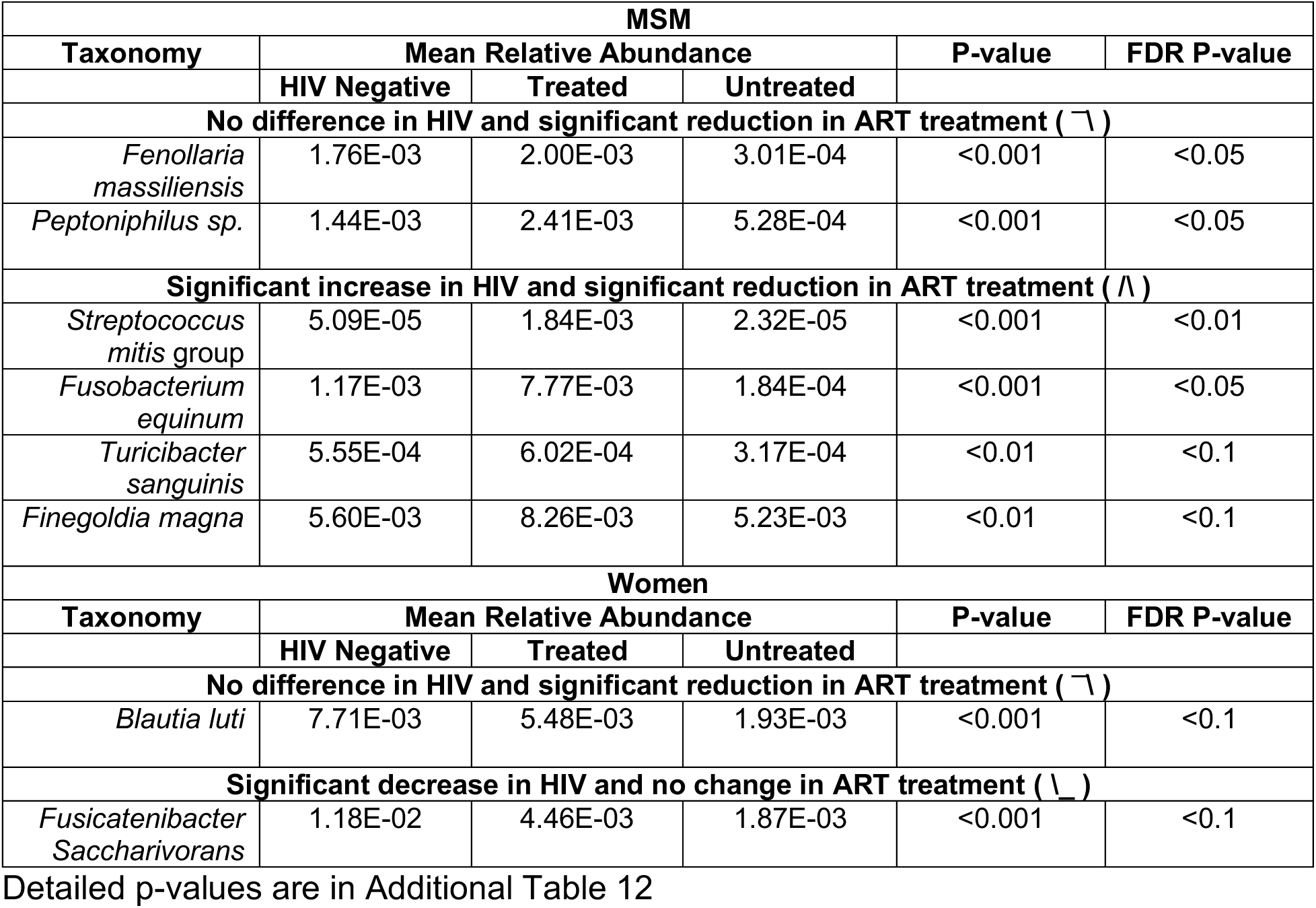
Significantly altered OTUs in HIV

PCoA analysis of women alone revealed that half of the HIV-positive women, both ART-treated and untreated, clustered with the HIV-negative women, with the other half clustering separately in the PCoA space using the “enterotyping” procedure (Figure 4B) (21). This clustering pattern was not significantly associated with CD4+ T cell count, viral load, CD4 nadir, CD4+CD38+HLA-DR+ and CD8+CD38+HLA-DR+ cells, antibiotic use in past 6 months, and engagement in RAI as assessed with either Kruskal-Wallis comparison of these factors between clusters or correlation with position of PC1 (primary axis of cluster division). Interestingly, these HIV positive women who were compositionally divergent from HIV negative women were characterized by microbiomes relatively high in Prevotella and low in Bacteroides (Additional Table 13), raising the intriguing possibility that this difference is driven by an unknown risk behavior-related factor, as the HIV-negative control women were not exhibiting any measured high-risk behaviors.

ART-naïve, HIV-positive women had significantly reduced alpha diversity compared to the seronegative controls (Kruskal-Wallis; p < 0.05; Figure 4C). Two OTUs were altered when comparing HIV-negative women; HIV-positive, ART-naïve women; and HIV-positive women on ART: *Fusicatenibacter saccharivorans* and *Blautia luti* (Table 3; Additional Table 12). *B. luti* had no significant difference in untreated HIV infection compared to HIV-negative controls and a significant decrease in treated HIV-infection compared to untreated infection. This suggests *B. luti* may be responding to ART and not HIV infection as a driver, however a failure to detect significant differences between HIV negative and ART naïve may have been a function of a low sample size in the ART naïve cohort. *F saccharivorans* had a significant decrease in relative abundance in untreated HIV-infection compared to HIV-negative individuals and no significant difference between untreated and treated HIV-positive individuals, suggesting an HIV-driven change that was not fully corrected by ART.

### Immune-microbiome correlations

To determine relationships between gut microbiome composition and markers of HIV disease progression/severity, we correlated the relative abundance of microbes in our HIV-positive, ART-naïve and ART-treated MSM with blood CD4 count, plasma HIV RNA (viral load), and percent of HLA-DR+CD38+ chronically activated CD4+ and CD8+ T cells (Figure 5). All of the significant correlations in the untreated population are negative, suggesting a loss of beneficial microbes with regard to inflammation especially when looking at the correlations with plasma viral load and CD8+HLA-DR+CD38+ percent. The CD4+HLA-DR+CD38+ correlations are more difficult to interpret: because activated CD4+ T cells are preferentially targeted in HIV infection, negative correlations may be due to a significant loss of these cells and not necessarily a reduction in activation.

**Figure 5.**
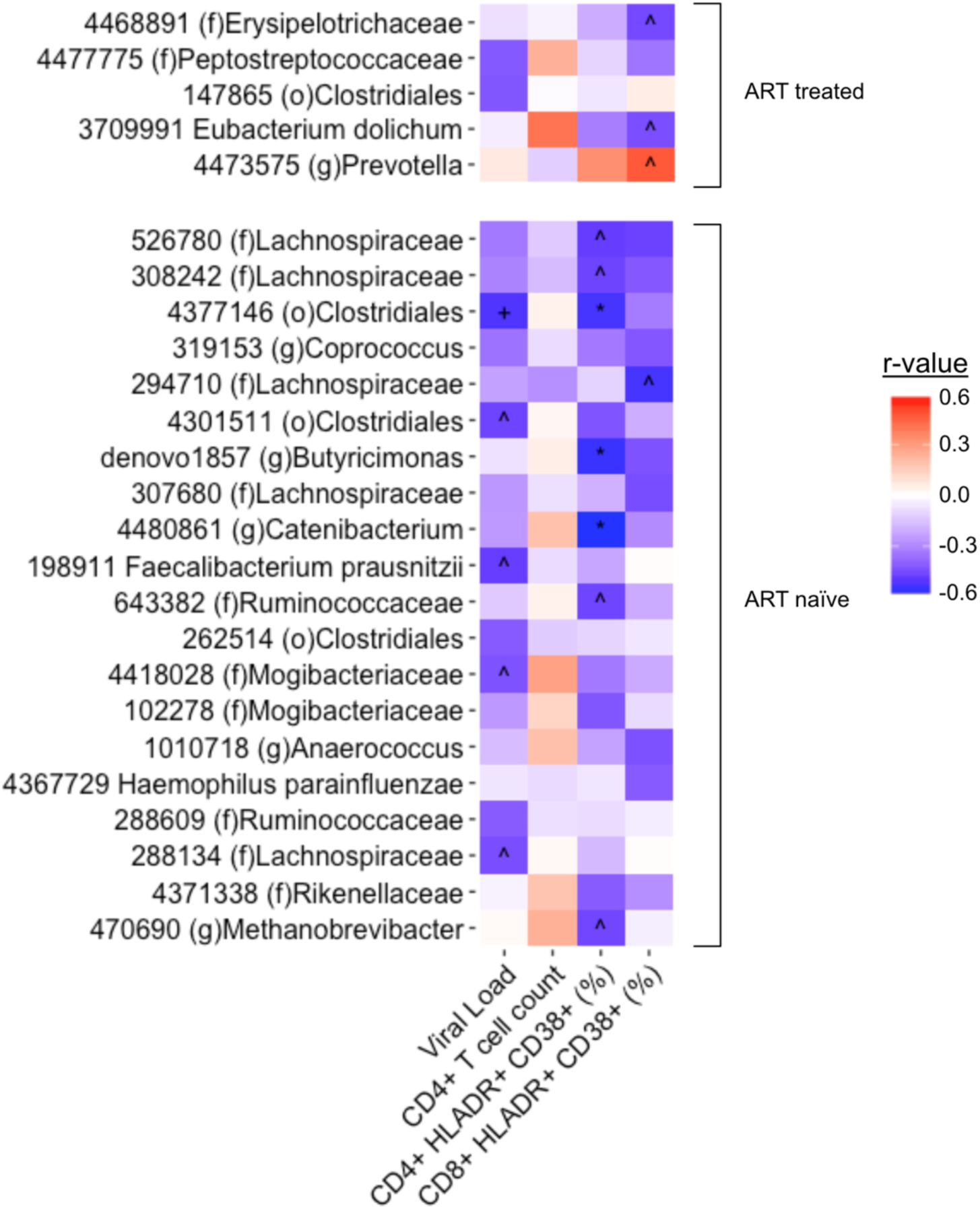
Microbiome immune correlations. Correlations in HIV-positive, ART-treated (top) and ART-naive (bottom) MSM between OTUs and serum measures of CD4+ T cell count, CD4+/CD8+ CD38+ HLA-DR+ cell percent, and viral load (All OTUs with one or more correlation r > |0.4| are displayed) (Spearman Rank test; FDR p-value: ^∧^ < 0.2, + < 0.1, * < 0.05).

In the ART-treated group, an OTU within the *Prevotella* genus, was positively correlated with CD8+HLA-DR+CD38+ percent. When compared against the NCBI 16S rRNA gene database using BLAST the OTU matched with 100% identity over 100% of the sequence to *Massiliprevotella massiliensis* strain Marseille-P2439 which is closely related to *Prevotella stercorea* (35). Additionally an OTU identified as *Eubacterium dolichum* was significantly negatively correlated with CD8+HLA-DR+CD38+ percent with a trending positive correlation with CD4+ T cell count, suggesting a potentially beneficial role.

### Longitudinal analysis of HIV-positive individuals before and after ART initiation

In addition to the cross-sectional cohort, we also studied 40 individuals longitudinally. HIV-negative (n=16) and HIV-positive (n=24) subjects were sampled 6-14 months apart. Individuals in the HIV-positive cohort were ART naïve at the first time point and then immediately initiated ART. The HIV-positive cohort had a significant increase in CD4+ T cell count (Wilcoxon signed-rank test; p < 0.05; Additional Figure 6) and decrease in viral load post-treatment with plasma HIV RNA counts between 0 and 40 (Wilcoxon signed-rank test; p < 0.001; Additional Figure 6).

HIV-positive individuals had significantly higher weighted but not unweighted beta diversity across time points compared to the seronegative controls, suggesting ART initiation results in significant alteration of the community proportions (UniFrac, Kruskal-Wallis Test, p < 0.05 (weighted), p = 0.11 (unweighted); Figure 6a). This result is also mirrored in abundance and binary Jaccard (Additional Figure 1D). There was no significant difference in the change in alpha diversity between the two time points when comparing HIV-positive individuals and HIV-negative controls (Figure 6B). However, two subjects had a significant reduction in observed OTUs and one subject had significant reduction in Shannon diversity as defined by being significant outliers from the rest of the data (Tietjen-Moore test, p < 0.05). While there was a 3-month antibiotic exclusion criteria for initial enrollment, subjects may have taken antibiotics between the two collected time points. However, there was no significant association between beta diversity or change in alpha diversity and antibiotic usage in the 6 months prior to the second sample collection (p > 0.05; Kruskal-Wallis test; Figure 6a,b). We also specifically tested whether the OTUs that had a significant reduction in relative abundance in ART-treated compared to untreated HIV infection in the cross-sectional analysis of MSM (Table 3), also were significantly reduced longitudinally after ART initiation. We found a significant reduction in the OTUs identified as *Turicibacter sanguinis* and *Streptococcus* sp. (*S. mitis* group) (Wilcoxon signed-rank test, p < 0.05; Figure 6c).

**Figure 6.**
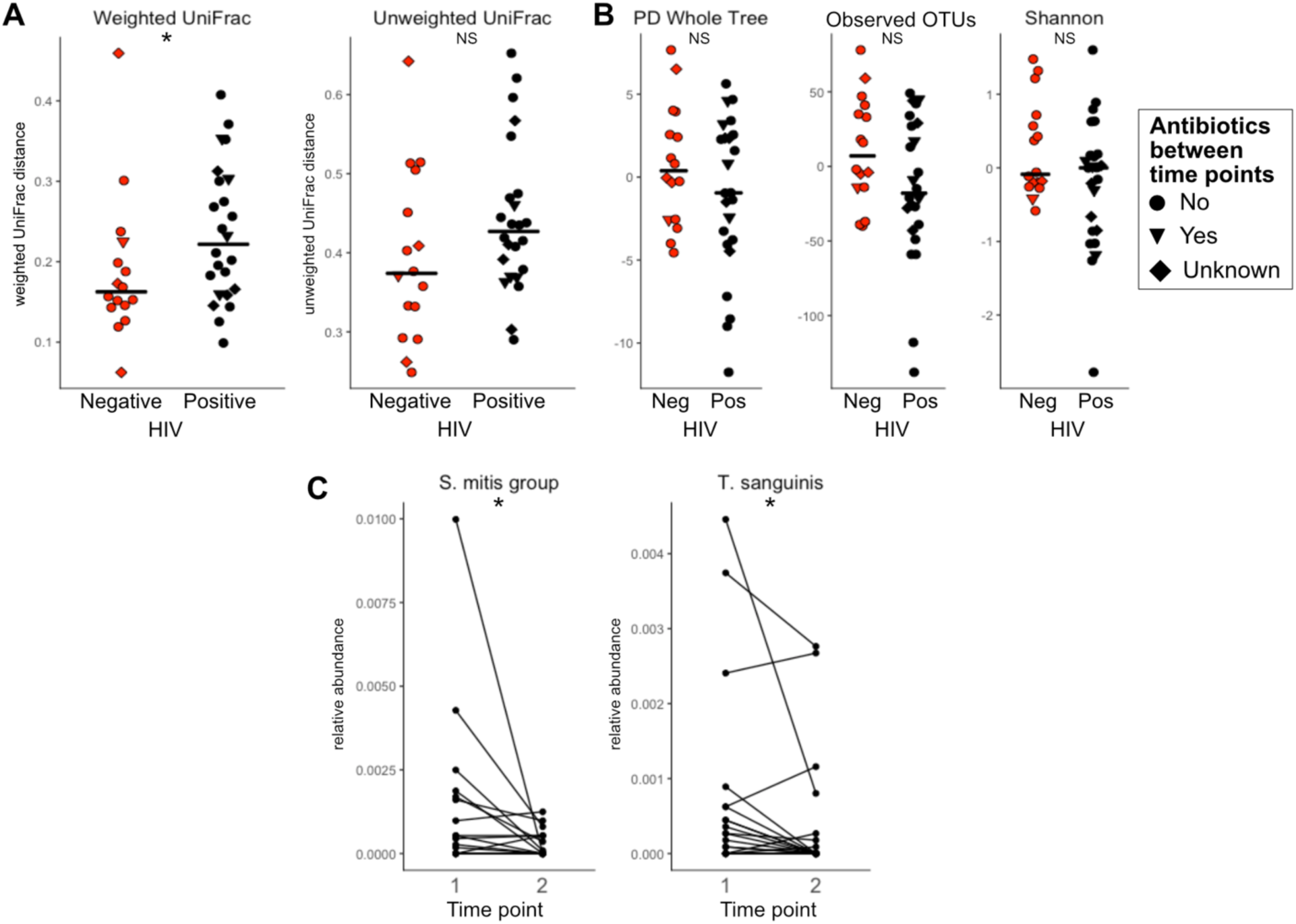
Longitudinal Analysis of ART. **A.** Beta diversity and **B.** change in alpha diversity of HIV negative individuals compared to HIV-positive pre- and post-ART initiation. **C.** Two taxa observed in the cross-sectional cohort as significantly reduced with HIV and ART are have significantly reduced relative abundance longitudinally (Wilcoxon signed-rank test; p-value: * < 0.05, NS = not significant).)

## Discussion

This study describes microbiome differences that occur in a US population infected with HIV and establishes the degree to which these differences may be driven by sexual practices, diet, ART, and HIV infection itself. Our data supports that Prevotella-rich microbiomes are associated with MSM and not independently with HIV infection status as has been previously reported (18). Surprisingly, given that Prevotella enterotypes have been associated with diets rich in carbohydrates and poor in animal products (21), we found MSM in our cohort to be consuming significantly less fiber and more red meat than non-MSM. This supports the conclusion of Noguera-Jullian et al. that dietary differences were not a likely driving factor of the Prevotella-rich MSM microbiome, especially given that both that study and ours found similar MSM-associated microbiome changes but different diet changes (18). Furthermore, when analyzing a subset of our cohort for which we had diet data with linear models that included kcal normalized amount of fiber or servings of red meat, we did not see a loss of significance of OTUs that differed between MSM and MSW/women. There were no microbiome associations with RAI engagement or frequency in MSM or women. Another potential driving factor of an altered microbiome in MSM could be a higher incidence of STDs, however we did not have enough data on STDs within our study population to examine this. Additionally, Noguera-Julian et al. did not find associations between Prevotella-richness and various STDs including human papillomavirus, hepatitis B, or hepatitis C infection (18). Taken together, we unfortunately still do not understand what drives a Prevotella-rich microbiomes in MSM.

Prevotella-rich microbiomes in MSM also display marked differences with Prevotella-rich microbiomes of non-MSM, suggesting potential differences in functional attributes. Compositional differences between Prevotella-rich microbiomes in different contexts may underlie contradictory reports in the literature regarding potential health implications.Some studies have suggested a health benefit with Prevotella enrichment such as improved glucose metabolism in the context of a high-fiber diet (22). Other studies suggest a detrimental effect indicated by associations with rheumatoid arthritis (24) and with insulin resistance (27). Experiments to test for causality have used the one commercially available strain of *P. copri* (DSM-18205/JCM 13464) and have shown this strain to have a wide range of effects: with detrimental effects such as greater insulin resistance and glucose intolerance (27) as well as aggravated DSS induced colitis (24). The exclusive use of a single strain in these experiments is not optimal. Consistent with strain level differences in *P. copri* detected in rheumatoid arthritis versus controls using shotgun metagenomic sequencing (24), we saw evidence of strain-level variation in Prevotella-rich MSM compared to non-MSM, with one OTU identified as *P. copri* (4475169) significantly more abundant in MSM and another OTU also identified as P *copri* (4482723) significantly less abundant (Figure 2B; Additional Table 6). Performing experiments with *P. copri* strains cultured directly from fecal samples of interest will be needed to understand a possible effect of strains on differing health phenotypes.

However, strong co-occurrence associations between *P. copri* – as well as *Prevotella* species in general – and other taxa suggests that such a strong emphasis on P. *copri* may not be warranted when trying to understand potential health implications of Prevotella-rich microbiomes. We observed many differences in composition and co-occurrence relationships in Prevotella-rich microbiomes of MSM compared to non-MSM, suggesting that there may be different sub-types of Prevotella-rich microbiomes. In a separate study, we have used *in vitro* stimulations with collections of whole intact bacterial cells isolated from feces, and found that MSM microbiomes induced more pro-inflammatory innate immune activation compared to non-MSM controls (2). Several OTUs more abundant in Prevotella-rich MSM compared to Prevotella-rich non-MSM, but not any OTUs in the *Prevotella* genus, correlated strongly with high CD4+ and CD8+ T cell activation in these *in vitro* stimulations (2). We have also previously used *in vitro* stimulation assays to show that one of these MSM and immune-activation correlated bacteria, *Holdamenella biformis* ATCC 27806 (formally *Eubacterium biforme*) but not *P. copri* DSM-18205 is far more pro-inflammatory than other gut commensals tested, inducing high TNF-α to IL-10 ratio in peripheral blood mononuclear cell (PBMC) stimulations (14). Further work will be needed to understand potential health implications of the MSM-associated gut microbiome and whether Prevotella itself or other co-occurring microbes are important drivers of functional phenotypes.

Since most microbiome studies aimed at understanding gut microbiome differences with HIV have not controlled for MSM behavior (14–16, 36–38), we still have an incomplete understanding of HIV-associated microbiome characteristics. Many of the species and genera that we have found to be increased in MSM compared to non-MSM have been previously reported to differ with HIV in studies not controlled for MSM risk behavior including *H. biformis* (14), *Prevotella* (14–17), *Catenibacterium* (14, 15), and *Desulfovibrio* (38). Additionally, we confirm a prior report of an increase in alpha diversity in MSM (18). The high number of MSM in our previous HIV-positive cohorts are likely to be the driver of our previously reported result of an increase in alpha diversity with HIV (38). This significant increase in alpha diversity in MSM challenges popular opinion that higher alpha diversity always equates to a healthier gut microbiome, especially with preliminary results suggesting the MSM microbiome may be more inflammatory (2). Unlike Noguera-Julian et al., we did not find a significant decrease in alpha diversity with HIV infection status in MSM (18). However, the cohort with the lowest alpha diversity in that study were immunologic non-responders to ART (i.e. individuals with poor CD4+ T cell recovery after ART) and our current cohort had few individuals who would fit this definition.

We confirm previous reports that there are relatively subtle differences in fecal microbiome composition with HIV infection in the absence of CD4+ T cell counts indicative of AIDS (18), or when analyzing populations dominated by heterosexual transmission (39). However, our analyses in MSM individuals and women did reveal some OTUs that associated with HIV and ART. Perhaps the most interesting pattern observed are taxa that increase compared to MSM controls with untreated HIV infection and decrease to levels indistinguishable from the controls with effective ART, as these may be microbes whose populations are particularly sensitive to CD4+ T cell control. All four OTUs that showed this trend were highly related to potential pathogens including *Turicibacter sanguinis* (40), *Fusobacterium equinum* (41), *Finegoldia magna* (42) and *Streptococcus* sp. (*S. mitis* group) (43). Interestingly, *F. magna*, *Streptococcus* and *Fusobacterium* have been found to co-occur in clinical samples (44). A recent study from our lab showed that, *in vitro*, microbiomes containing higher levels of *T. sanguinis* induced more elevated frequencies of HLA-DR+CD38+CD4+ T cells, a marker of chronic activation, when whole microbiome bacterial isolates were cultured with PBMCs (2), suggesting a potential mechanistic relationship between *T. sanguinis* and T cell activation in the gut with untreated HIV infection. For the *Streptococcus mitis* group related OTU, the 16S rRNA gene sequence was 100% identical in the sequenced V4 region to 7 different type species with high variation in genome content (45). This highlights the need for isolation of bacteria directly from the study population and/or metagenomic sequencing to elucidate strain-specific changes in the microbiome and for follow-up experiments to establish functional implications.

Although our analysis of women showed stronger microbiome differences with HIV infection, with a proportion of HIV positive women clustering apart from HIV negative at the community level, these results should be interpreted with caution since our control cohort was not recruited as women with high-risk behaviors. Notably, the enrichment of *Prevotella* and depletion of *Bacteroides* characterizing the HIV-positive women that diverged from HIV negative women at the community level suggests the same unknown risk behavior may be a driving factor. Despite the variation in microbiome compositions of HIV-positive women, we observed significant differences in an OTU identified in the *Blautia* genus with HIV infection. This is consistent with a previously reported increase of *Blautia* in HIV-positive, ART-naïve individuals compared to HIV-negative controls in a heterosexual transmission dominated cohort recruited in Sweden (46).

In our longitudinal cohort, we observed more change in the microbiomes of HIV-positive individuals pre- and post-ART initiation compared to the HIV-negative controls over similar time intervals, suggesting that microbiome compositional changes after ART initiation are greater than the baseline stochasticity of the healthy human microbiome.Microbiome changes observed in our cross-sectional cohort were validated in our longitudinal cohort; specifically *Turicibacter sanguinis*, and *Streptococcus* sp. (*S. mitis* group) are significantly decreased with ART in both analyses, adding to our confidence in these results. ART associated changes are likely driven by a combination of beneficial changes caused by restoration of immune function and viral suppression and potentially detrimental changes driven by the ART drugs themselves. The changes driven by the ART drugs themselves likely vary based on drug regimen. For instance in a study conducted in Mexico City, Mexico, unique microbiome changes were observed in individuals on non-nucleoside reverse transcriptase inhibitors compared to ritonavir-boosted protease inhibitors with the same backbone of nucleoside reverse transcriptase inhibitors (26). However, the individuals in our cohort are on a wide variety of different ART drug combinations and we did not have enough power to determine ART-class associated changes in the microbiome.

## Conclusions

The gut microbiome in HIV is a potential contributor to infection, disease progression, and development of non-infectious HIV-associated diseases. However, developing an understanding of changes occurring with HIV infection has been complicated by large differences in the gut microbiomes of MSM, a population disproportionately affected by HIV in the US. We observed significant microbiome changes in HIV-negative MSM populations both in the prevalence of Prevotella-rich microbiomes as well as a different microbiome composition compared to Prevotella-rich non-MSM. Potential health implications and driving factors of Prevotella-rich microbiomes have been discussed in other health contexts including metabolic and autoimmune diseases, and understanding context-specific differences in Prevotella-rich microbiomes may help deconvolute contradicting results.

When controlling for MSM, we observed HIV- and ART-associated changes in the microbiome in both MSM and in women, some of which were confirmed in a longitudinal cohort of HIV-positive individuals initiating ART, showing an increase in potential pathogens in untreated infection that have the potential to contribute to inflammatory phenotypes. Lastly, both our correlational analysis as well as the greater body of literature suggest a potential influence of the microbiome composition in HIV on the inflammatory state of the HIV-infected gut. This understanding will help to guide efforts to investigate the functional implications of these differences to ultimately target the microbiome to improve the health of this population.

## Methods

### Subject recruitment

Subjects were residents of the Denver, Colorado metropolitan area and were recruited and studied at the Clinical Translational Research Center of the University of Colorado Hospital. Three cohorts were prospectively recruited based on HIV status. 1) HIV-1 infection untreated: Individuals with a positive antibody or PCR test at least 6 months prior to enrollment and either ART drug-naïve (defined as < 10 days of ART treatment at any time prior to entry), or previously on ART but off treatment for the previous 6 months prior to screening. 2) Chronic HIV-1 infection on long-term ART: ART for ≥12 months with a minimum of three ART drugs prior to study entry and <50 copies HIV RNA/mL within 30 days prior to study entry and no plasma HIV-1 RNA ≥50 copies/mL in the past 6 months. 3) Healthy controls: HIV negative individuals both high and low-risk for contracting HIV. High-risk for HIV infection was defined as in a prior study of a candidate HIV vaccine: 1) a history of unprotected anal intercourse with one or more male or male-to-female transgender partners; 2) anal intercourse with two or more male or male-to-female transgender partners; or 3) being in a sexual relationship with a person who has been diagnosed with HIV (47). Individuals who reported taking antibiotics within 3 months of sample collection were excluded from the study. The fecal microbiome data from 50 of the 217 subjects were described in our previous publications (14, 38)

### Longitudinal Cohort

A subset of HIV-positive, untreated patients returned for a second visit and sample collection after 6-14 months of ART. A control cohort of HIV-negative participants also provided two samples 6-14 months apart. Analysis of microbiome change (beta diversity and delta alpha diversity) over time revealed no significant effect of variation in time between samples within a 6-14 month window (Spearman-rank test; P> 0.05); therefore we did not account for time between samples in our analysis.

### DNA Extraction and Sequencing

Stool samples were collected on sterile swabs by the patient within 24 hours prior to their clinic visit and stored at −80°C. DNA was extracted using the standard DNeasy PowerSoil Kit protocol (Qiagen). Extracted DNA was PCR amplified with barcoded primers targeting the V4 region of 16S rRNA gene according to the Earth Microbiome Project 16S Illumina Amplicon protocol with the 515F:806R primer constructs (48). Control sterile swab samples that had undergone the same DNA extraction and PCR amplification procedures were also processed. Each PCR product was quantified using PicoGreen (Invitrogen), and equal amounts (ng) of DNA from each sample were pooled and cleaned using the UltraClean PCR Clean-Up Kit (MoBio). Sequences were generated on six runs on a MiSeq personal sequencer (Illumina, San Diego, CA).

### Sequence Data Analysis

Raw sequences were demultiplexed using QIIME 1.9.1 (49). DADA2 (50) was used for error correcting and unique sequence identification. During DADA2 processing all sequences were truncated to 112 bp. Unique sequences were then binned into 99% OTUs using sortmeRNA (closed reference) (51) and UCLUST (open reference)(52) in QIIME 1.9.1 and taxonomy was assigned using the GreenGenes database 18_3 (53). Samples were rarefied at 11,218 reads. Beta diversity metrics were calculated using Weighted and Unweighted UniFrac (54, 55) and binary and abundance Jaccard (56). Alpha diversity metrics were calculated using phylogenetic diversity whole tree (30), observed OTUs, and Shannon using QIIME. Co-occurrence network modeling was performed using an in-house python script utilizing sparCC (57) implemented as fast sparCC (https://github.com/shafferm/fast_sparCC) in the programming language, Python.

### Diet Information

Food frequency questionnaires were collected using Diet History Questionnaire II (58). Diet composition was processed using the Diet*Calc software and the dhq2.database.092914 database (59). All reported values are based on USDA nutrition guidelines. Reported dietary levels were both normalized per 1000 kcal and unnormalized.

### Behavioral Assessment

A behavioral questionnaire to assess sexual preference and risk behaviors within the past year was administered to study participants either at the time of fecal sample collection or retroactively. Data on sexual preference was used to assess MSM status in all individuals but data on practices such as frequency of RAI were only used for individuals who filled out the questionnaire within a year of fecal sample collection.

### Immunologic Assays

Whole blood was collected in BD Vacutainer tubes containing EDTA. 50μL of blood was surface stained for CD4 counts with anti-CD3, anti-CD4, anti-CD45 (BD Biosciences TriTest), and CD8 counts with anti-CD3, anti-CD8, anti-CD45 (BD Biosciences Tritest). Red blood cells (RBCs) were lysed with FACS Lysis Solution (BD) and fixed with 1% formaldehyde. 50μL of blood was surface stained for CD38 and HLA-DR counts on CD4 and CD8 cells using anti-CD4/CD8, anti-CD38, anti-CD3, and anti-HLA- DR (BD Biosciences Multitest). RBCs were lysed (BD) and fixed with 1% formaldehyde. Analysis was performed on a BD FACSCALIBUR using Multiset software.

### Statistical Analysis

Non-parametric statistical tests were performed in the R software package (v 3.3.2; http://www.r-project.org). All correlations were performed using Spearman correlation. Analysis of variance was calculated using Kruskal-Wallis test. Paired longitudinal analysis was calculated using Wilcoxon signed-rank test. P values less than 0.05 were considered significant. The Benjamini-Hochberg Procedure was used to correct for multiple tests, when applicable, (FDR p): FDR p values less than 0.1 were considered significant. The R package, vegan, was used to perform the adonis test at 10000 permutations (60).

## Declarations

### Ethics Approval and consent to participate

Informed consent was obtained from each subject, and the study protocol was approved by the Colorado Multiple Institution Review Board (CoMIRB 09-0898 13-2986 14-1595, 15-1692).

### Consent for publication

Not applicable

### Availability of data and material

The datasets analyzed during this study are available in the European Nucleotide Archive repository [PRJEB28485] (https://www.ebi.ac.uk/ena/data/view/PRJEB28485).

### Competing interests

The authors declare that they have no competing interests.

### Funding

This work was funded by R01 DK108366, R01 DK104047, U01 HL098996, and U01 HL121816. This work also was supported by additional MicroGrant funding and other general resources from NIH/NCATS Colorado CTSA Grant Number UL1 TR002535. AJS Armstrong was supported by T32 AI052066 and M Shaffer was supported by 2 T15 LM009451-06. Contents are the authors’ sole responsibility and do not necessarily represent official NIH views.

### Authors’ contributions

AJSA analyzed and interpreted all data. AJSA and CAL wrote the manuscript. CG and SF recruited subjects and collected samples. NMN prepared and ran sequencing. AJSA and MS wrote computational resources. CPN and JMS performed immune assays. AF, TC, BEP, and CAL procured funding. All authors read and approved the final manuscript.

## Acknowledgements

We would like to thank our study participants for contributing their samples and time to this study. Thanks to Rob Knight for use of compute resources.

## Additional Figures

**Additional Figure 1.**
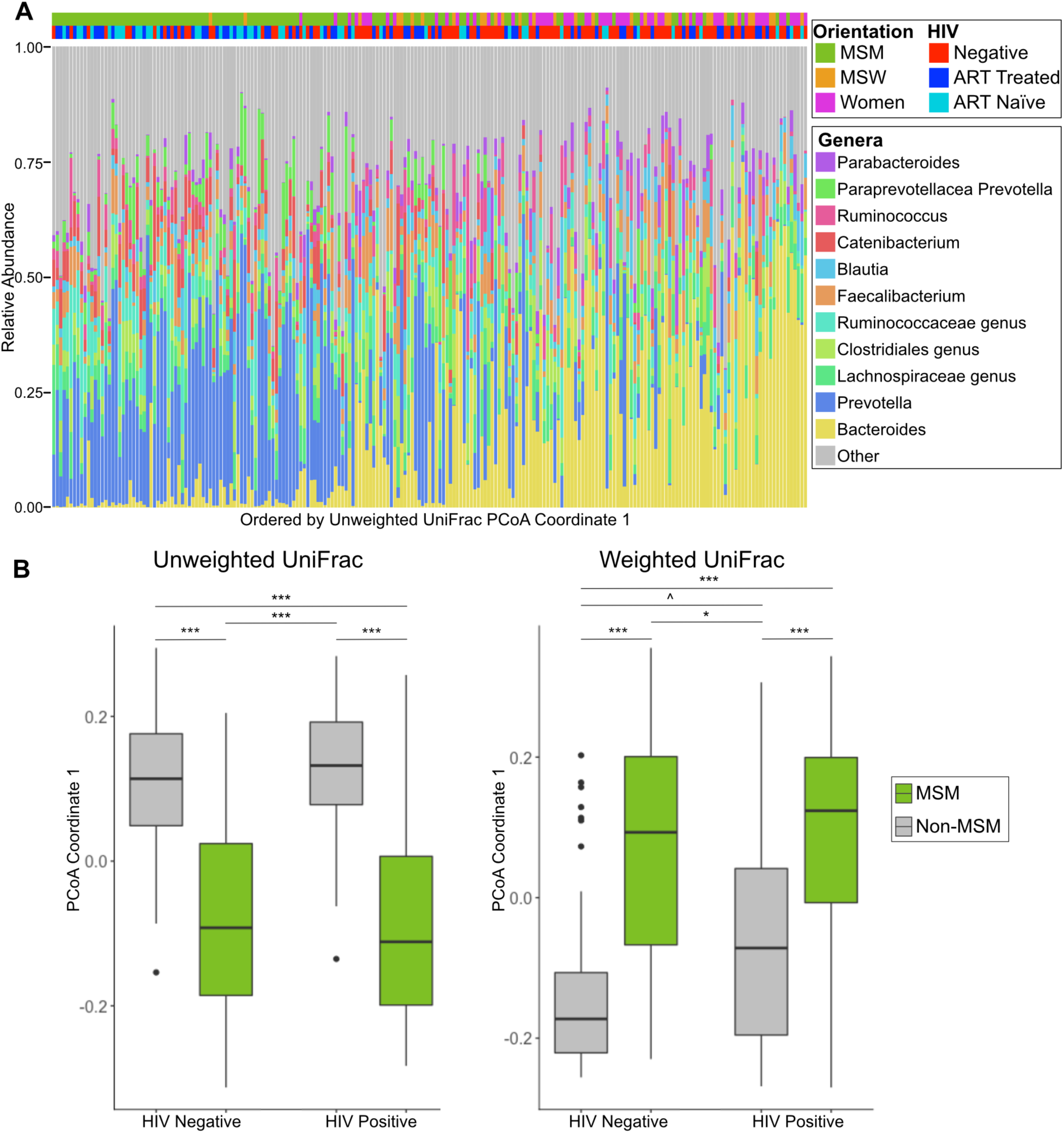
Principal coordinate analysis for taxonomy and clustering. **A.** Genus level taxa bar chart order by unweighted UniFrac PCoA coordinate 1. (Bottom) Genera with mean relative less than 2% abundance are binned together into the category “Other”. Each column represents one individual. (Top) samples are marked with HIV status and orientation. Each column corresponds to the genus plot below. Samples are ordered by the coordinate on the weighted UniFrac principle coordinate 1. **B.** Orientation and HIV “clustering” along principal coordinate 1 in unweighted (left) and weighted (right) UniFrac. (Kruskal-Wallis test, p<0.001; Dunn’s Post Hoc Test, FDR P-value: ^∧^ < 0.1, * < 0.05, ** < 0.01, *** < 0.001)

**Additional Figure 2.**
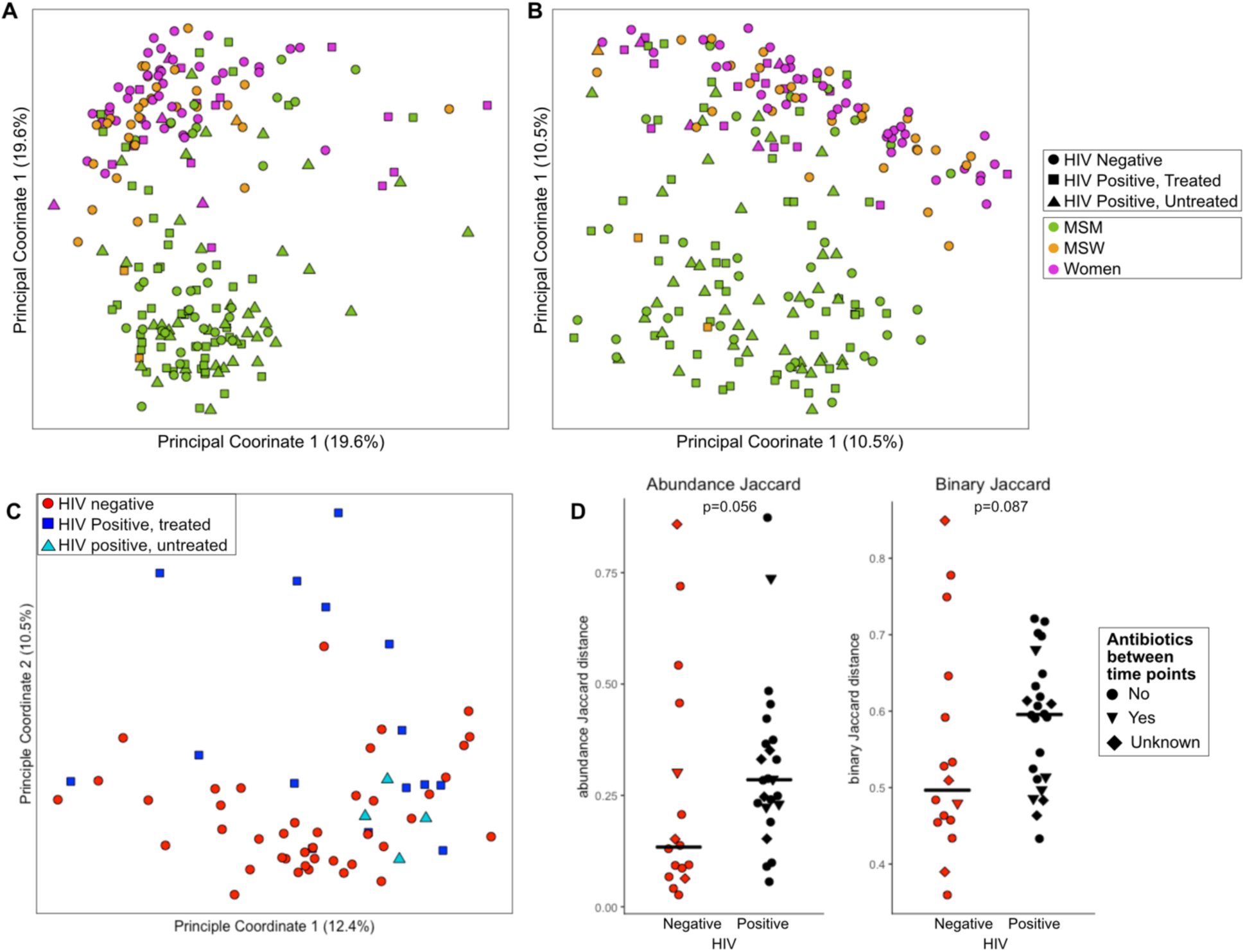
Abundance and Binary Jaccard Analysis. **A.** Abundance and **B.** binary Jaccard with points colored by orientation and shaped by HIV status in all study samples. **C.** Abundance Jaccard of women only; points colored by HIV status. **D.** Abundance (left) and binary (right) Jaccard of longitudinal samples comparing HIV negative individuals to HIV-positive pre- and post-ART initiation (Kruskal-Wallis test).

**Additional Figure 3.**
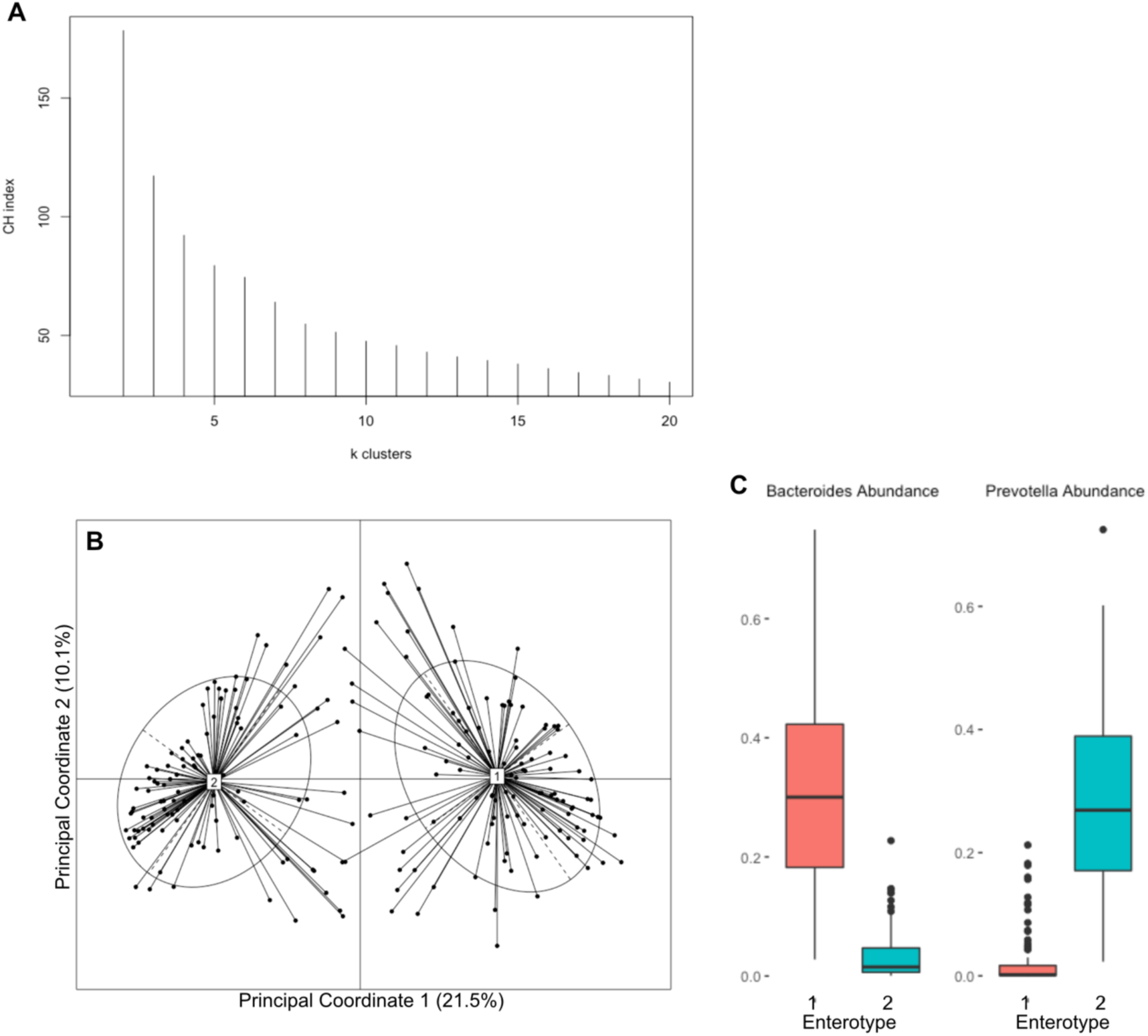
Samples are best divided into two enterotypes, which are primarily defined by Bacteroides or Prevotella richness. **A.** Silhouette analysis shows that two clusters is the most efficient way to divide the data. **B.** PCoA of Jensen-Shannon divergence distance. The center of each enterotype/cluster is marked with a line to each member of the cluster. Centers were calculated using Partitioning around medoids (PAM). **C.** The relative abundance of the two most abundant taxa in the clusters, Prevotella and Bacteroides. Each enterotype is marked by a Prevotella-rich/Bacteroides-poor or Bacteroides-rich/Prevotella-poor composition.

**Additional Figure 4.**
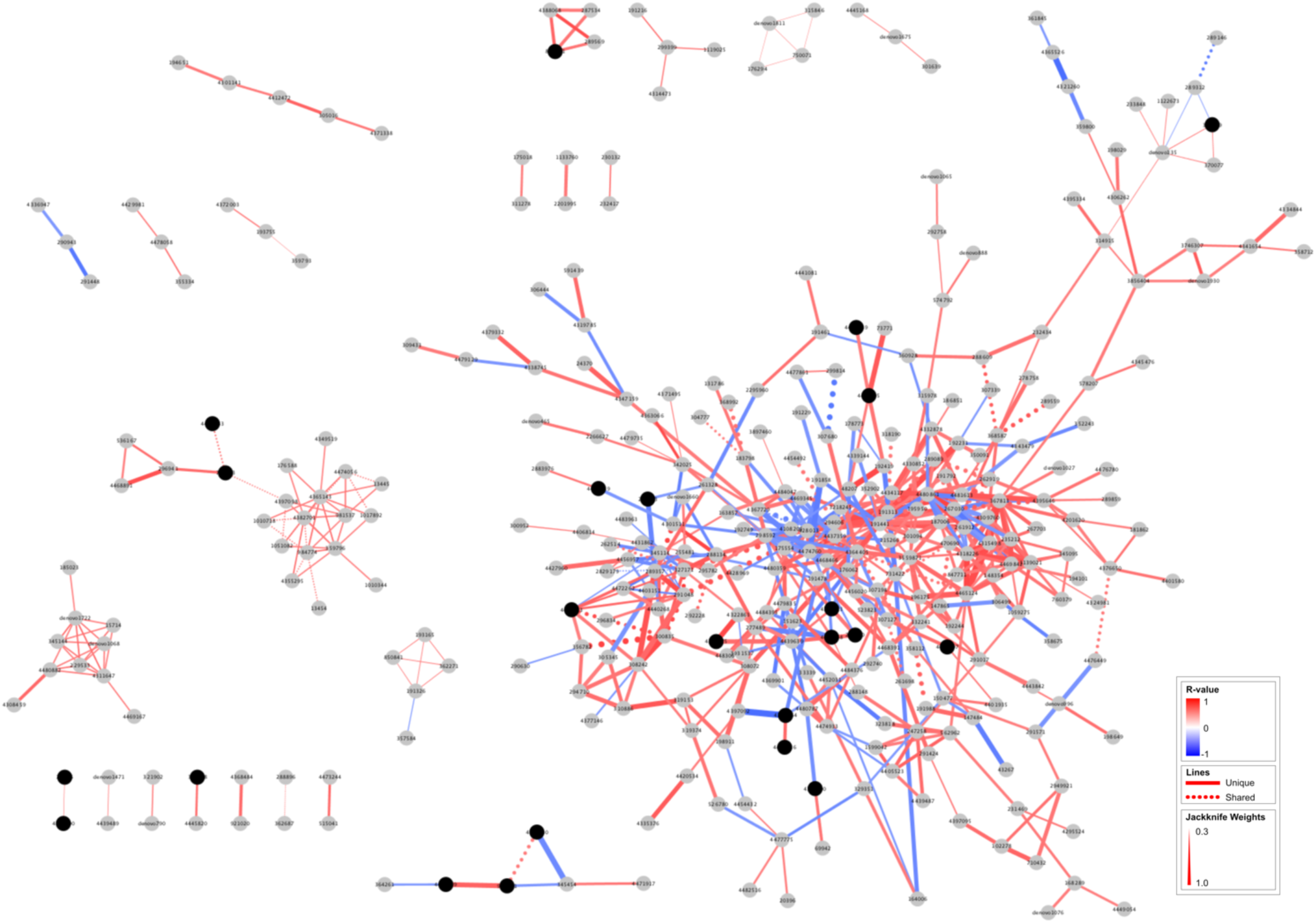
Co-occurrence network of HIV-negative, Prevotella-rich MSM. Correlation network of the 345 OTUs in the 25 HIV-negative, Prevotella-rich MSM individuals calculated using SparCC. The value of the correlation is colored on the network edges with negative R-values in blue and positive in red. Only correlations greater than |0.5| are included in the network. Analysis was jackknifed 100 times on random subsets of 9 individuals. The edge weights from the jackknife are represented by line weight, with the thickest lines corresponding to correlations identified in all jackknife subsets. The network edges on this graph are compared to a network calculated using the same methods with a subset of 9 individuals that are HIV-negative, Prevotella-rich non-MSM. The edges that are shared between the MSM and non-MSM networks are dotted and the edges unique to the MSM network are solid. The OTUs identified in the Prevotella genus are colored black.

**Additional Figure 5.**
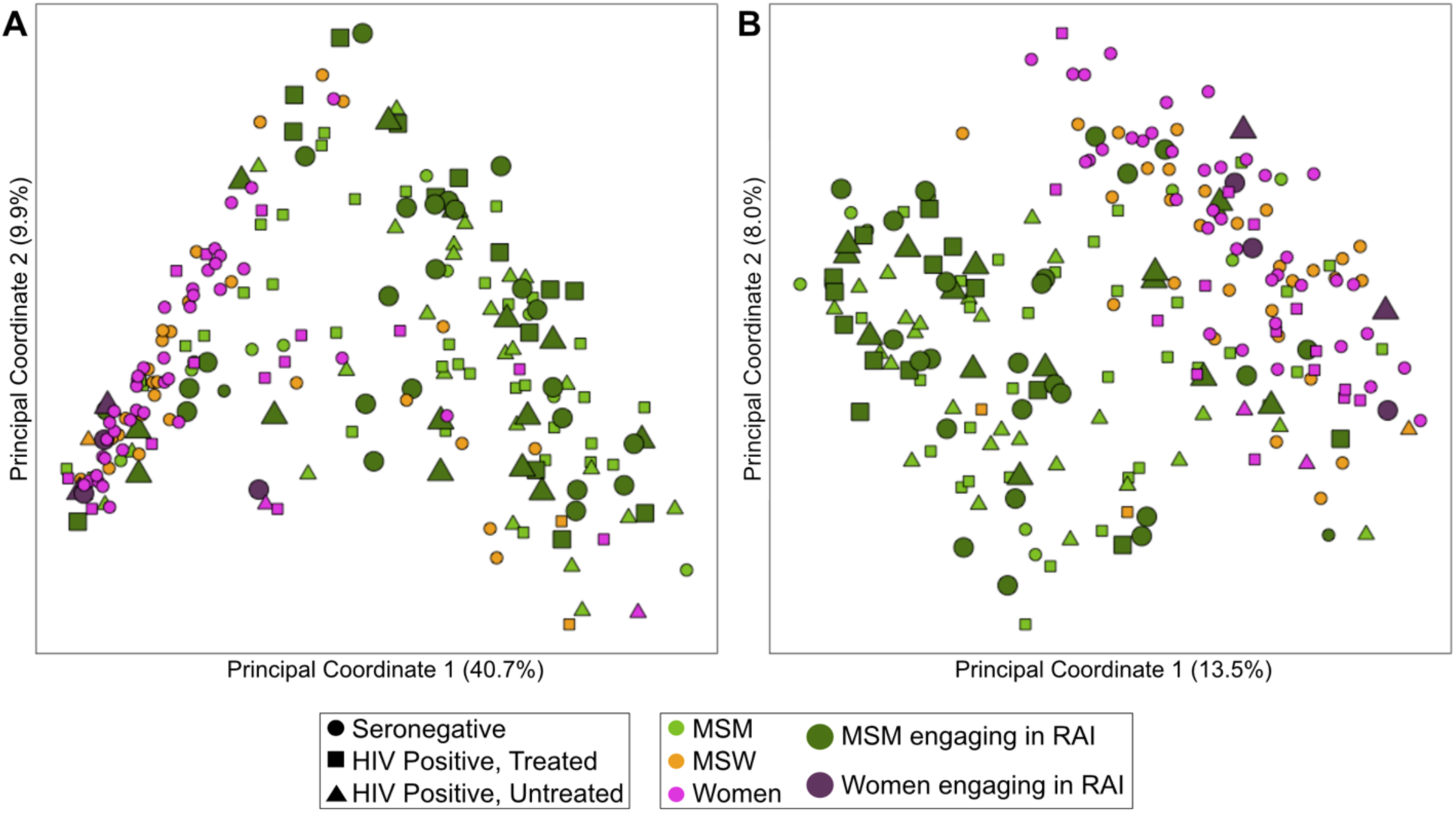
Receptive anal intercourse does not correlate with PCoA space. **A.** Weighted UniFrac PCoA and **B.** Unweighted UniFrac with points colored by orientation and shaped by HIV status. Women who reported engaging in RAI are colored dark purple and are larger in size than the other points. This plot highlights that women who engage in RAI cluster apart from the MSM with the other Bacteroides-rich women.

**Additional Figure 6.**
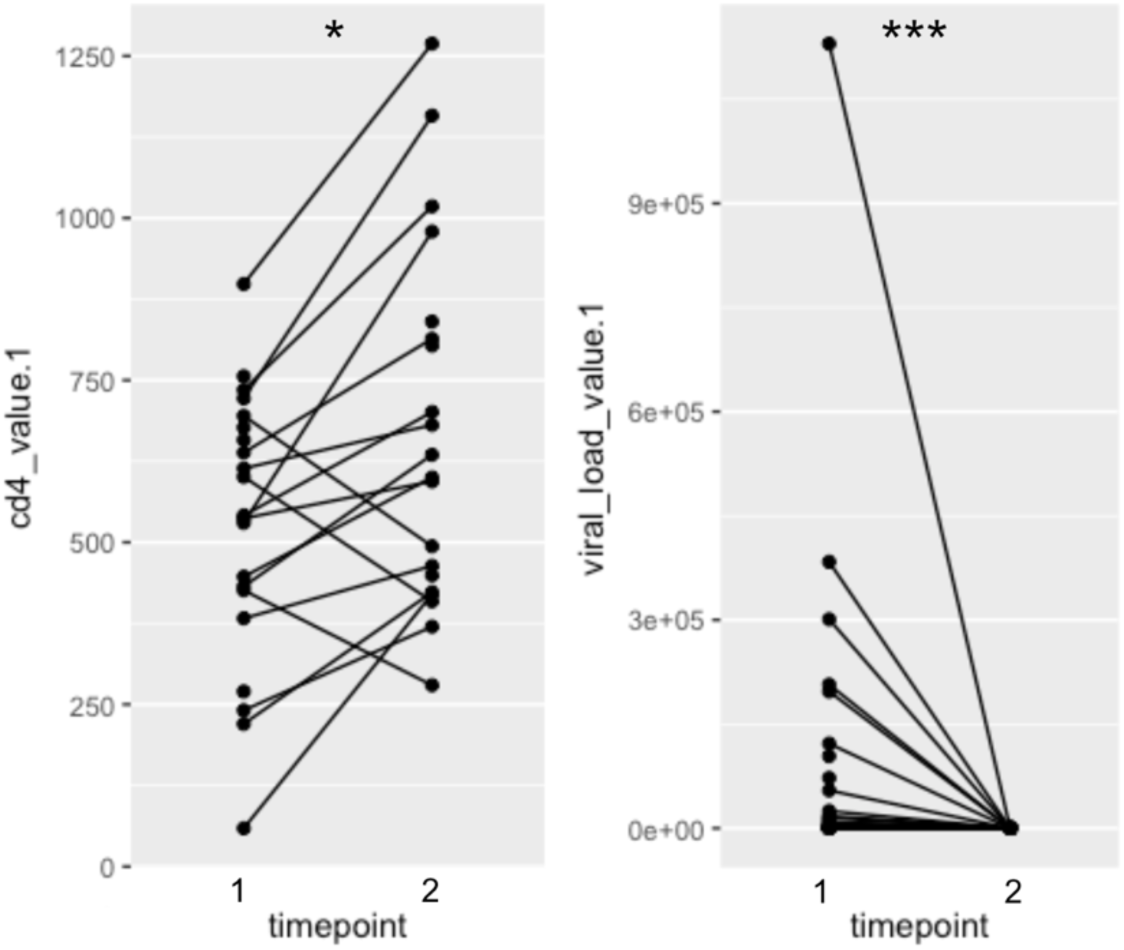
HIV-positive individuals improve CD4+ T cell count and viral load after ART initiation. (Wilcoxon rank-sum; P-value: * < 0.05, *** < 0.001).

## Additional Table Legends

**Additional Table 1. (xlsx) Table of statistical analyses of differences in parameters across groups in Table 1.**

**Additional Table 2. (xlsx) Table of statistical analyses of differences in parameters across groups in Table 2.**

**Additional Table 3. (xlsx) Adonis test results for contribution of HIV, MSM and ART status to differences in weighted and unweighted UniFrac distance matrices.**Results are presented for the full cohort as well as a sensitivity analysis that excluded HIV-positive men with unknown orientation.

**Additional Table 4. (xlsx) Table of significantly different OTUs between all (HIV-negative and positive) MSM and non-MSM.** OTUs significantly increased with MSM highlighted in orange OTUs significantly decreased in MSM highlighted in green (Kruskal-Wallis test).

**Additional Table 5. (xlsx) Fishers exact test for enterotypes.org fit of MSM compared to non-MSM to reference enterotypes.**

**Additional Table 6. (xlsx) Table of significantly different OTUs between only HIV-negative MSM and non-MSM.** OTUs significantly increased with MSM highlighted in orange OTUs significantly decreased in MSM highlighted in green (Kruskal-Wallis test).

**Additional Table 7. (xlsx) Edge Table for HIV-negative, Prevotella-rich MSM network displayed in Additional Figure 4.**

**Additional Table 8. (xlsx) Edge Table for HIV-negative, Prevotella-rich non-MSM network referenced in in Additional Figure 4.**

**Additional Table 9. (xlsx) Full list of diet differences in MSM compared to non-MSM normalized by 1000kcal.** Diet components significantly increased with MSM highlighted in orange OTUs significantly decreased in MSM highlighted in green (Kruskal-Wallis test).

**Additional Table 10. (xlsx) Full list of diet differences in MSM compared to non-MSM un-normalized data.** Diet components significantly increased with MSM highlighted in orange OTUs significantly decreased in MSM highlighted in green (Kruskal-Wallis test).

**Additional Table 11. (xlsx) Linear modeling of diet and MSM results analysis to determine effect of diet differences on OTUs that differed between MSM and non-MSM/females.**

**Additional Table 12. (xlsx) The Dunn’s test statistic for OTUs identified to differ between MSM who are HIV-negative, HIV-positive untreated, and HIV-positive ART using the Kruskal-Wallis test in Table 3.**

**Additional Table 13. (xlsx) Significantly different genera between the two clusters in the women-only comparison of HIV-positive and HIV-negative individuals.**

## List of abbreviations

ART: Antiretroviral Therapy
FTM: Female to Male
HIV: Human Immunodeficiency Virus
MSM: Men who have Sex with Men
MSW: Men who have Sex with Women
OTU: Operational Taxonomic Unit
PBMC: Peripheral Blood Mononuclear Cell
PCoA: Principal Coordinates Analysis
PD: Phylogenetic Diversity
RAI: Receptive Anal Intercourse

## References

1. UNAIDS. 2017. Latest global and regional statistics on the status of the AIDS epidemic. http://www.unaids.org/sites/default/files/mediaasset/UNAIDSFactSheeten.pdf. Accessed

2. Neff CP, Krueger O, Xiong K, Arif S, Nusbacher N, Schneider JM, Cunningham AW, Armstrong A, Li S, McCarter MD, Campbell TB, Lozupone CA, Palmer BE. 2018. Fecal Microbiota Composition Drives Immune Activation in HIV-infected Individuals. EBioMedicine 30:192–202.

3. Taha TE, Hoover DR, Dallabetta GA, Kumwenda NI, Mtimavalye LA, Yang LP, Liomba GN, Broadhead RL, Chiphangwi JD, Miotti PG. 1998. Bacterial vaginosis and disturbances of vaginal flora: association with increased acquisition of HIV. AIDS 12:1699–706.

4. Burgener A, McGowan I, Klatt NR. 2015. HIV and mucosal barrier interactions: consequences for transmission and pathogenesis. Curr Opin Immunol 36:22–30.

5. CDC. 2017. HIV in the United States: At A Glance. In: PREVENTION, C. F. D. C. A. (ed.). Accessed

6. Somsouk M, Estes JD, Deleage C, Dunham RM, Albright R, Inadomi JM, Martin JN, Deeks SG, McCune JM, Hunt PW. 2015. Gut epithelial barrier and systemic inflammation during chronic HIV infection. AIDS (London, England) 29:43–51.

7. Dieffenbach CW, Fauci AS. 2011. Thirty years of HIV and AIDS: future challenges and opportunities. Annals of internal medicine 154:766–71.

8. Montessori V, Press N, Harris M, Akagi L, Montaner JSG. 2004. Adverse effects of antiretroviral therapy for HIV infection. CMAJ: Canadian Medical Association journal = journal de l’Association medicale canadienne 170:229–38.

9. Kelley CF, Kraft CS, de Man TJ, Duphare C, Lee HW, Yang J, Easley KA, Tharp GK, Mulligan MJ, Sullivan PS, Bosinger SE, Amara RR. 2017. The rectal mucosa and condomless receptive anal intercourse in HIV-negative MSM: implications for HIV transmission and prevention. Mucosal Immunol 10:996–1007.

10. Vijay-Kumar M, Aitken JD, Carvalho FA, Cullender TC, Mwangi S, Srinivasan S, Sitaraman SV, Knight R, Ley RE, Gewirtz AT. 2010. Metabolic syndrome and altered gut microbiota in mice lacking Toll-like receptor 5. Science (New York, NY) 328:228–31.

11. Koeth RA, Wang Z, Levison BS, Buffa JA, Org E, Sheehy BT, Britt EB, Fu X, Wu Y, Li L, Smith JD, DiDonato JA, Chen J, Li H, Wu GD, Lewis JD, Warrier M, Brown JM, Krauss RM, Tang WHW, Bushman FD, Lusis AJ, Hazen SL. 2013. Intestinal microbiota metabolism of L-carnitine, a nutrient in red meat, promotes atherosclerosis. Nat Med 19:576–85.

12. Honda K, Littman DR. 2012. The Microbiome in Infectious Disease and Inflammation. Annu Rev Immunol 30:759–795.

13. Li SX, Armstrong AJS, Neff CP, Shaffer M, Lozupone CA, Palmer BE. 2016. Complexities of gut microbiome dysbiosis in the context of HIV infection and antiretroviral therapy. Clinical pharmacology and therapeutics 00:1–12.

14. Lozupone CA, Li M, Campbell TB, Flores SC, Linderman D, Gebert MJ, Knight R, Fontenot AP, Palmer BE. 2013. Alterations in the gut microbiota associated with HIV-1 infection. Cell host & microbe 14:329–39.

15. Mutlu EA, Keshavarzian A, Losurdo J, Swanson G, Siewe B, Forsyth C, French A, Demarais P, Sun Y, Koenig L, Cox S, Engen P, Chakradeo P, Abbasi R, Gorenz A, Burns C, Landay A. 2014. A compositional look at the human gastrointestinal microbiome and immune activation parameters in HIV infected subjects. PLoS Path 10:e1003829.

16. Dillon SM, Lee EJ, Kotter CV, Austin GL, Dong Z, Hecht DK, Gianella S, Siewe B, Smith DM, Landay AL, Robertson CE, Frank DN, Wilson CC. 2014. An altered intestinal mucosal microbiome in HIV-1 infection is associated with mucosal and systemic immune activation and endotoxemia. Mucosal Immunology 7:983–94.

17. Vázquez-Castellanos JF, Serrano-Villar S, Latorre A, Artacho A, Ferrús ML, Madrid N, Vallejo A, Sainz T, Martínez-Botas J, Ferrando-Martínez S, Vera M, Dronda F, Leal M, Del Romero J, Moreno S, Estrada V, Gosalbes MJ, Moya A. 2015. Altered metabolism of gut microbiota contributes to chronic immune activation in HIV-infected individuals. Mucosal immunology 8:760–72.

18. Noguera-Julian M, Rocafort M, Guillén Y, Rivera J, Casadellá M, Nowak P, Hildebrand F, Zeller G, Parera M, Bellido R, Rodríguez C, Carrillo J, Mothe B, Coll J, Bravo I, Estany C, Herrero C, Saz J, Sirera G, Torrela A, Navarro J, Crespo M, Brander C, Negredo E, Blanco J, Guarner F, Calle ML, Bork P, Sönnerborg A, Clotet B, Paredes R. 2016. Gut Microbiota Linked to Sexual Preference and HIV Infection. EBioMedicine 5:135–46.

19. Ley RE. 2016. Gut microbiota in 2015: Prevotella in the gut: choose carefully. Nature Publishing Group 13:69–70.

20. Arumugam M, Raes J, Pelletier E, Le Paslier D, Yamada T, Mende DR, Fernandes GR, Tap J, Bruls T, Batto J-M, Bertalan M, Borruel N, Casellas F, Fernandez L, Gautier L, Hansen T, Hattori M, Hayashi T, Kleerebezem M, Kurokawa K, Leclerc M, Levenez F, Manichanh C, Nielsen HB, Nielsen T, Pons N, Poulain J, Qin J, Sicheritz-Ponten T, Tims S, Torrents D, Ugarte E, Zoetendal EG, Wang J, Guarner F, Pedersen O, de Vos WM, Brunak S, Doré J, Antolín M, Artiguenave F, Blottiere HM, Almeida M, Brechot C, Cara C, Chervaux C, Cultrone A, Delorme C, Denariaz G, Dervyn R, et al. 2011. Enterotypes of the human gut microbiome. Nature 473:174–80.

21. Wu GD, Chen J, Hoffmann C, Bittinger K, Chen Y-Y, Keilbaugh SA, Bewtra M, Knights D, Walters WA, Knight R, Sinha R, Gilroy E, Gupta K, Baldassano R, Nessel L, Li H, Bushman FD, Lewis JD. 2011. Linking long-term dietary patterns with gut microbial enterotypes. Science (New York, NY) 334:105–8.

22. Kovatcheva-Datchary P, Nilsson A, Akrami R, Lee Ying S, De Vadder F, Arora T, Hallen A, Martens E, Björck I, Bäckhed F. 2015. Dietary Fiber-Induced Improvement in Glucose Metabolism Is Associated with Increased Abundance of Prevotella. Cell Metabolism 22:971–982.

23. Albenberg LG, Wu GD. 2014. Diet and the Intestinal Microbiome: Associations, Functions, and Implications for Health and Disease. Gastroenterology 146:1564–1572.

24. Scher JU, Sczesnak A, Longman RS, Segata N, Ubeda C, Bielski C, Rostron T, Cerundolo V, Pamer EG, Abramson SB, Huttenhower C, Littman DR. 2013. Expansion of intestinal Prevotella copri correlates with enhanced susceptibility to arthritis. eLife 2:e01202.

25. Zhang H, DiBaise JK, Zuccolo A, Kudrna D, Braidotti M, Yu Y, Parameswaran P, Crowell MD, Wing R, Rittmann BE, Krajmalnik-Brown R. 2009. Human gut microbiota in obesity and after gastric bypass. Proc Natl Acad Sci U S A 106:2365–70.

26. Pinto-Cardoso S, Lozupone C, Briceno O, Alva-Hernandez S, Tellez N, Adriana A, Murakami-Ogasawara A, Reyes-Teran G. 2017. Fecal Bacterial Communities in treated HIV infected individuals on two antiretroviral regimens. Sci Rep 7:43741.

27. Pedersen HK, Gudmundsdottir V, Nielsen HB, Hyotylainen T, Nielsen T, Jensen BA, Forslund K, Hildebrand F, Prifti E, Falony G, Le Chatelier E, Levenez F, Dore J, Mattila I, Plichta DR, Poho P, Hellgren LI, Arumugam M, Sunagawa S, VieiraSilva S, Jorgensen T, Holm JB, Trost K, Meta HITC, Kristiansen K, Brix S, Raes J, Wang J, Hansen T, Bork P, Brunak S, Oresic M, Ehrlich SD, Pedersen O. 2016. Human gut microbes impact host serum metabolome and insulin sensitivity. Nature 535:376–81.

28. Gianella S, Strain MC, Rought SE, Vargas MV, Little SJ, Richman DD, Spina CA, Smith DM. 2012. Associations between virologic and immunologic dynamics in blood and in the male genital tract. J Virol 86:1307–15.

29. Palmer CD, Tomassilli J, Sirignano M, Romero-Tejeda M, Arnold KB, Che D, Lauffenburger DA, Jost S, Allen T, Mayer KH, Altfeld M. 2014. Enhanced immune activation linked to endotoxemia in HIV-1 seronegative MSM. AIDS 28:2162–6.

30. Faith DP. 1994. Phylogenetic pattern and the quantification of organismal biodiversity. Philos Trans R Soc Lond B Biol Sci 345:45–58.

31. Costea PI, Hildebrand F, Arumugam M, Backhed F, Blaser MJ, Bushman FD, de Vos WM, Ehrlich SD, Fraser CM, Hattori M, Huttenhower C, Jeffery IB, Knights D, Lewis JD, Ley RE, Ochman H, O’Toole PW, Quince C, Relman DA, Shanahan F, Sunagawa S, Wang J, Weinstock GM, Wu GD, Zeller G, Zhao L, Raes J, Knight R, Bork P. 2018. Enterotypes in the landscape of gut microbial community composition. Nat Microbiol 3:8–16.

32. Gevers D, Knight R, Petrosino JF, Huang K, McGuire AL, Birren BW, Nelson KE, White O, Methe BA, Huttenhower C. 2012. The Human Microbiome Project: a community resource for the healthy human microbiome. PLoS Biol 10:e1001377.

33. Qin J, Li R, Raes J, Arumugam M, Burgdorf KS, Manichanh C, Nielsen T, Pons N, Levenez F, Yamada T, Mende DR, Li J, Xu J, Li S, Li D, Cao J, Wang B, Liang H, Zheng H, Xie Y, Tap J, Lepage P, Bertalan M, Batto J-M, Hansen T, Le Paslier D, Linneberg A, Nielsen HB, Pelletier E, Renault P, Sicheritz-Ponten T, Turner K, Zhu H, Yu C, Li S, Jian M, Zhou Y, Li Y, Zhang X, Li S, Qin N, Yang H, Wang J, Brunak S, Doré J, Guarner F, Kristiansen K, Pedersen O, Parkhill J, Weissenbach J, et al. 2010. A human gut microbial gene catalogue established by metagenomic sequencing. Nature 464:59–65.

34. Volpe GE, Ward H, Mwamburi M, Dinh D, Bhalchandra S, Wanke C, Kane AV. 2014. Associations of cocaine use and HIV infection with the intestinal microbiota, microbial translocation, and inflammation. Journal of studies on alcohol and drugs 75:347–57.

35. Ndongo S, Cadoret F, Dubourg G, Delerce J, Fournier PE, Raoult D, Lagier JC. 2017. ‘Collinsella phocaeensis’ sp. nov., ‘Clostridium merdae’ sp. nov., ‘Sutterella massiliensis’ sp. nov., ‘Sutturella timonensis’ sp. nov., ‘Enorma phocaeensis’ sp. nov., ‘Mailhella massiliensis’ gen. nov., sp. nov., ‘Mordavella massiliensis’ gen. nov., sp. nov. and ‘Massiliprevotella massiliensis’ gen. nov., sp. nov., 9 new species isolated from fresh stool samples of healthy French patients. New Microbes New Infect 17:89–95.

36. Nowak P, Troseid M, Avershina E, Barqasho B, Neogi U, Holm K, Hov JR, Noyan K, Vesterbacka J, Svard J, Rudi K, Sonnerborg A. 2015. Gut microbiota diversity predicts immune status in HIV-1 infection. AIDS 29:000–000.

37. McHardy IH, Li X, Tong M, Ruegger P, Jacobs J, Borneman J, Anton P, Braun J. 2013. HIV Infection is associated with compositional and functional shifts in the rectal mucosal microbiota. Microbiome 1:26.

38. Lozupone CA, Rhodes ME, Neff CP, Fontenot AP, Campbell TB, Palmer BE. 2014. HIV-induced alteration in gut microbiota. Gut Microbes 5:562–570.

39. Monaco CL, Gootenberg DB, Zhao G, Handley SA, Ghebremichael MS, Lim ES, Lankowski A, Baldridge MT, Wilen CB, Flagg M, Norman JM, Keller BC, Luevano JM, Wang D, Boum Y, Martin JN, Hunt PW, Bangsberg DR, Siedner MJ, Kwon DS, Virgin HW. 2016. Altered Virome and Bacterial Microbiome in Human Immunodeficiency Virus-Associated Acquired Immunodeficiency Syndrome. Cell Host Microbe 19:311–22.

40. Bosshard PP, Zbinden R, Altwegg M. 2002. Turicibacter sanguinis gen. nov., sp. nov., a novel anaerobic, Gram-positive bacterium. Int J Syst Evol Microbiol 52:1263–6.

41. Tadepalli S, Stewart GC, Nagaraja TG, Narayanan SK. 2008. Human Fusobacterium necrophorum strains have a leukotoxin gene and exhibit leukotoxic activity. J Med Microbiol 57:225–31.

42. Goto T, Yamashita A, Hirakawa H, Matsutani M, Todo K, Ohshima K, Toh H, Miyamoto K, Kuhara S, Hattori M, Shimizu T, Akimoto S. 2008. Complete genome sequence of Finegoldia magna, an anaerobic opportunistic pathogen. DNA Res 15:39–47.

43. Mitchell J. 2011. Streptococcus mitis: walking the line between commensalism and pathogenesis. Mol Oral Microbiol 26:89–98.

44. Lee S, Roh KH, Kim CK, Yong D, Choi JY, Lee JW, Lee K, Chong Y. 2008. A case of necrotizing fasciitis due to Streptococcus agalactiae, Arcanobacterium haemolyticum, and Finegoldia magna in a dog-bitten patient with diabetes. Korean J Lab Med 28:191–5.

45. Kawamura Y, Hou XG, Todome Y, Sultana F, Hirose K, Shu SE, Ezaki T, Ohkuni H. 1998. Streptococcus peroris sp. nov. and Streptococcus infantis sp. nov., new members of the Streptococcus mitis group, isolated from human clinical specimens. Int J Syst Bacteriol 48 Pt 3:921–7.

46. Vesterbacka J, Rivera J, Noyan K, Parera M, Neogi U, Calle M, Paredes R, Sonnerborg A, Noguera-Julian M, Nowak P. 2017. Richer gut microbiota with distinct metabolic profile in HIV infected Elite Controllers. Sci Rep 7:6269.

47. Hammer SM, Sobieszczyk ME, Janes H, Karuna ST, Mulligan MJ, Grove D, Koblin BA, Buchbinder SP, Keefer MC, Tomaras GD, Frahm N, Hural J, Anude C, Graham BS, Enama ME, Adams E, DeJesus E, Novak RM, Frank I, Bentley C, Ramirez S, Fu R, Koup RA, Mascola JR, Nabel GJ, Montefiori DC, Kublin J, McElrath MJ, Corey L, Gilbert PB, Team HS. 2013. Efficacy trial of a DNA/rAd5 HIV-1 preventive vaccine. N Engl J Med 369:2083–92.

48. Thompson LR, Sanders JG, McDonald D, Amir A, Ladau J, Locey KJ, Prill RJ, Tripathi A, Gibbons SM, Ackermann G, Navas-Molina JA, Janssen S, Kopylova E, Vazquez-Baeza Y, Gonzalez A, Morton JT, Mirarab S, Zech Xu Z, Jiang L, Haroon MF, Kanbar J, Zhu Q, Jin Song S, Kosciolek T, Bokulich NA, Lefler J, Brislawn CJ, Humphrey G, Owens SM, Hampton-Marcell J, Berg-Lyons D, McKenzie V, Fierer N, Fuhrman JA, Clauset A, Stevens RL, Shade A, Pollard KS, Goodwin KD, Jansson JK, Gilbert JA, Knight R, Earth Microbiome Project C. 2017. A communal catalogue reveals Earth’s multiscale microbial diversity. Nature 551:457–463.

49. Caporaso JG, Kuczynski J, Stombaugh J, Bittinger K, Bushman FD, Costello EK, Fierer N, Peña AG, Goodrich JK, Gordon JI, Huttley GA, Kelley ST, Knights D, Koenig JE, Ley RE, Lozupone CA, McDonald D, Muegge BD, Pirrung M, Reeder J, Sevinsky JR, Turnbaugh PJ, Walters WA, Widmann J, Yatsunenko T, Zaneveld J, Knight R. 2010. QIIME allows analysis of high-throughput community sequencing data. Nat Methods 7:335–6.

50. Callahan BJ, McMurdie PJ, Rosen MJ, Han AW, Johnson AJ, Holmes SP. 2016. DADA2: High-resolution sample inference from Illumina amplicon data. Nat Methods 13:581–3.

51. Kopylova E, Noe L, Touzet H. 2012. SortMeRNA: fast and accurate filtering of ribosomal RNAs in metatranscriptomic data. Bioinformatics 28:3211–7.

52. Edgar RC. 2010. Search and clustering orders of magnitude faster than BLAST. Bioinformatics 26:2460–1.

53. McDonald D, Price MN, Goodrich J, Nawrocki EP, DeSantis TZ, Probst A, Andersen GL, Knight R, Hugenholtz P. 2012. An improved Greengenes taxonomy with explicit ranks for ecological and evolutionary analyses of bacteria and archaea. The ISME journal 6:610–8.

54. Lozupone C, Knight R. 2005. UniFrac: a new phylogenetic method for comparing microbial communities. Appl Environ Microbiol 71:8228–35.

55. Lozupone CA, Hamady M, Kelley ST, Knight R. 2007. Quantitative and qualitative beta diversity measures lead to different insights into factors that structure microbial communities. Appl Environ Microbiol 73:1576–85.

56. Jaccard P. 1912. The Distribution of the Flora in the Alpine Zone.1. New Phytol 11:37–50.

57. Friedman J, Alm EJ. 2012. Inferring correlation networks from genomic survey data. PLoS Comput Biol 8:e1002687.

58. National Institues of Health EaGRP, Nactional Cancer Institute. 2010. Diet History Questionnaire, Verson 2.0.

59. National Cancer Institute EaGRP. 2012. Diet*Calc Analysis Program, Version 1. 5.0..

60. Jari Oksanen FGB, Michael Friendly, Roeland Kindt, Pierre Legendre, Dan McGlinn, Peter R. Minchin, R. B. O’Hara, Gavin L. Simpson, Peter Solymos, M. Henry H. Stevens, Eduard Szoecs and Helene Wagner. 2017. vegan: Community Ecology Package., vR package version 2.4-4. https://CRAN.R-project.org/package=vegan.

